# Compensatory mutations potentiate constructive neutral evolution by gene duplication

**DOI:** 10.1101/2024.02.12.579783

**Authors:** Philippe C Després, Alexandre K Dubé, Jordan Grenier, Marie-Ève Picard, Rong Shi, Christian R Landry

## Abstract

Protein functions generally depend on their assembly into complexes. During evolution, some complexes have transitioned from homomers encoded by a single gene to heteromers encoded by duplicate genes. This transition could occur without adaptive evolution through intermolecular compensatory mutations. Here, we experimentally duplicate and evolve an homodimeric enzyme to examine if and how this could happen. We identify hundreds of deleterious mutations that inactivate individual homodimers but produce functional enzymes when co-expressed as duplicated proteins that heterodimerize. The structure of one such heteromer reveals how both losses of function are buffered through the introduction of asymmetry in the complex that allows them to subfunctionalize. Constructive neutral evolution can thus occur by gene duplication followed by only one deleterious mutation per duplicate.

**One sentence summary:** Compensatory deleterious mutations entangle gene duplicates

## Main text

Some protein complexes with equivalent functions exist as homomers in some species and as heteromers of paralogous proteins in others (*1*, *2*). While the transitions from one form to the other may be favored by selection due to gain of novel functions or the optimization of ancestral ones, it has also been proposed that non-adaptive mechanisms could sometimes be sufficient to explain this complexification (*3–5*). One of the main models for the maintenance of gene duplicates proposes that complementary loss-of-function (LOF) mutations after gene duplication make both gene copies essential to maintain the ancestral function (*6*). Complementation between LOF mutations could occur at the physical level when paralogs heterodimerize through *trans* epistatic effects. Such compensatory mutations would result in heteromeric complexes that maintain the ancestral function and that at the same time make duplicated genes co-dependent (Figure 1a). Even if the resulting heteromers have slightly lower activity than the ancestral complex, this transition could occur neutrally or near neutrally, as the duplication itself could mask the impact of the first deleterious mutation and because metabolic networks can usually buffer significant variation in fluxes without affecting fitness (*7*).

**Figure 1.**
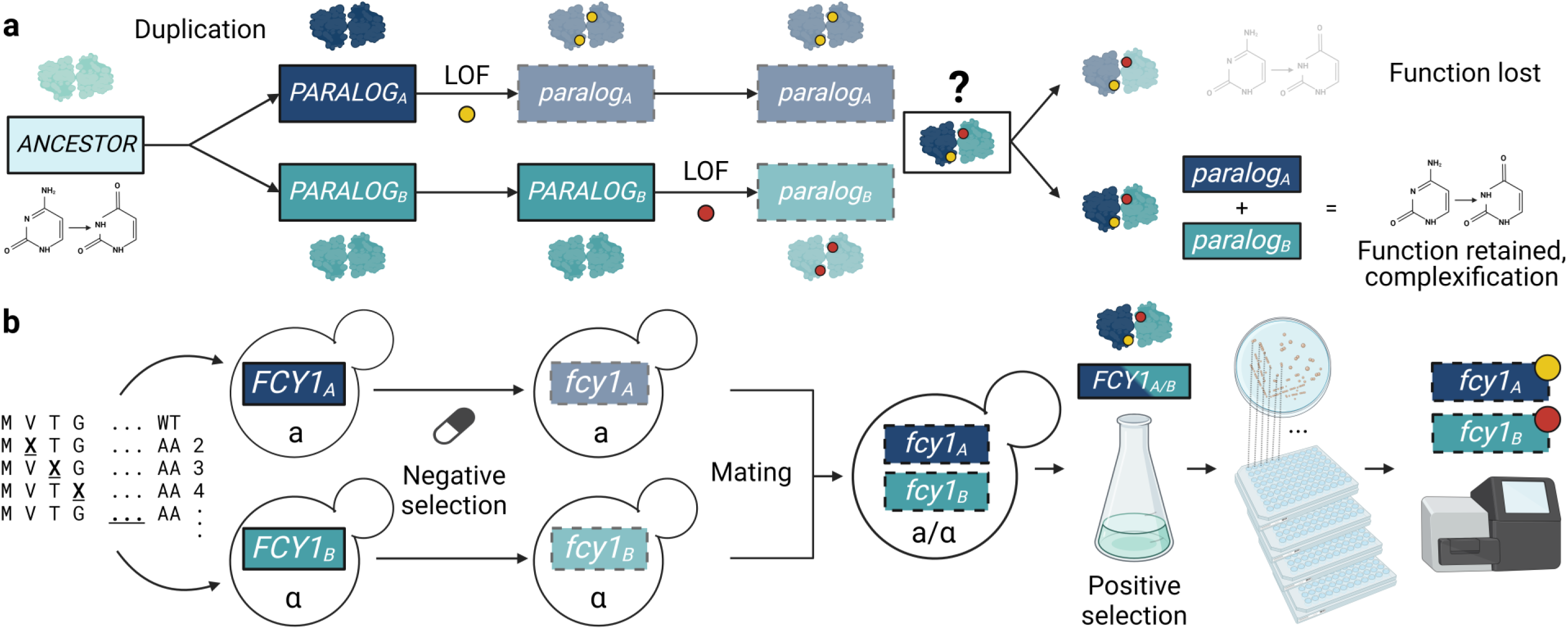
Compensatory mutations for the replacement of a homodimer by a heterodimer of paralogs. **a)**. After a duplication event, independent loss of function (LOF) events in each of the young paralogs abolishes the activity of both homomeric protein complexes. If there is i ntermolecular compensation of sufficient magnitude between the two inactive paralogous subunits, then the heteromer can maintain some level of activity and prevent the loss of ancestral function but with increased complexity. The two paralogs and their mutations mutually compensate for each other’s LOF. **b)** Experimental system to detect positive *trans* epistasis after an experimentally simulated duplication. Systematic mutagenesis libraries of *FCY1* integrated at the endogenous locus were generated in both yeast mating types a and α. The mutant libraries were enriched for LOF variants by negative selection (5 -FC media). The resulting pools of mutants were then crossed with one another and exposed to positive selection (cytosine media) to enrich for restored complex function via *trans* epistasis. From these pools, 2,304 single colonies were isolated for one-by-one *FCY1* genotyping using high-throughput sequencing. Figure 1 was created using Biorender.

To make heteromerization through compensatory deleterious mutations a likely event, there must exist a relatively large set of LOF mutations with the potential for strong intermolecular positive epistasis (epistasis in *trans*). Furthermore, the transition must occur through a limited number of mutations if it is to occur before one gene copy is lost and the system goes back to the ancestral single gene state. Currently, little is known about whether or not this requirement can be met, only that for some protein complexes this scenario could explain their gain in complexity over time (*8*) and the fact that many duplicated gene pairs are physically co-dependent (*9*).

To measure these parameters in an extant protein complex, we used the yeast cytosine deaminase (Fcy1, also known as yCD), a homodimeric enzyme, to simulate a homomer to heteromer evolutionary transition. Our measurements of the *in vitro* K_D_ of Fcy1 suggest it is an obligate homodimer, with over 90% of protein chains existing as part of a complex *in vivo* (Figure S1). Fcy1 activity potentiates the toxicity of the antifungal 5-FC (*10*), allowing for selection of nonfunctional protein variants in a pooled competition assay (*11*). Conversely, this complex also catalyzes the deamination of cytosine into uracil, restoring the growth of auxotrophic strains in media lacking uracil but supplemented with cytosine (cytosine media). Our previous work (*11*) revealed that 5-FC resistance is strongly negatively correlated with canonical Fcy1 activity through single amino acid mutations so that resistance to the drug is inevitably linked to strong loss of function. Using these two opposed selection pressures and the yeast sexual cycle, we designed an experiment to detect and characterize positive *trans* epistasis between pairs of nonfunctional gene copies thereby simulating a gene duplication followed by two successive parallel LOF events (Figure 1b). Compensation between duplicates would result in the phenotypic complementation of uracil auxotrophy as observed for the ancestral protein complex.

We built systematic single amino acid mutant libraries of *FCY1* at the endogenous locus in both yeast mating types and enriched for LOF mutations using negative selection. Fitness effects in these conditions were highly consistent between replicates and with our previous assay (Figure S2-5). We mated these pools to generate diploid strains bearing two *FCY1* gene copies (*FCY1*_A_ and *FCY1*_B_), mimicking genotypes with two gene copies as would occur after gene duplication in a haploid genome. The resulting libraries covered almost all pairwise LOF combinations. We exposed this diverse set of Fcy1_A_-Fcy1_B_ pairs to positive selection to enrich and isolate those that compensated each others’ LOF through the formation of heteromeric complexes. We identified 207 different functional gene pairs among at least 613 independent crossing events as assessed by sequencing (Figure S6-7). Almost half of the unique combinations were observed independently multiple times, representing a high-confidence set of positive *trans* epistatic interactions (Figure 2a). The diversity of pairs of mutations observed suggests there exists more compensating pairs in the sequence space (Figure S8). Strong compensatory effects between non-functional duplicates due to a single amino acid substitution are thus not restricted to a few unique mutation combinations but are instead relatively frequent.

**Figure 2.**
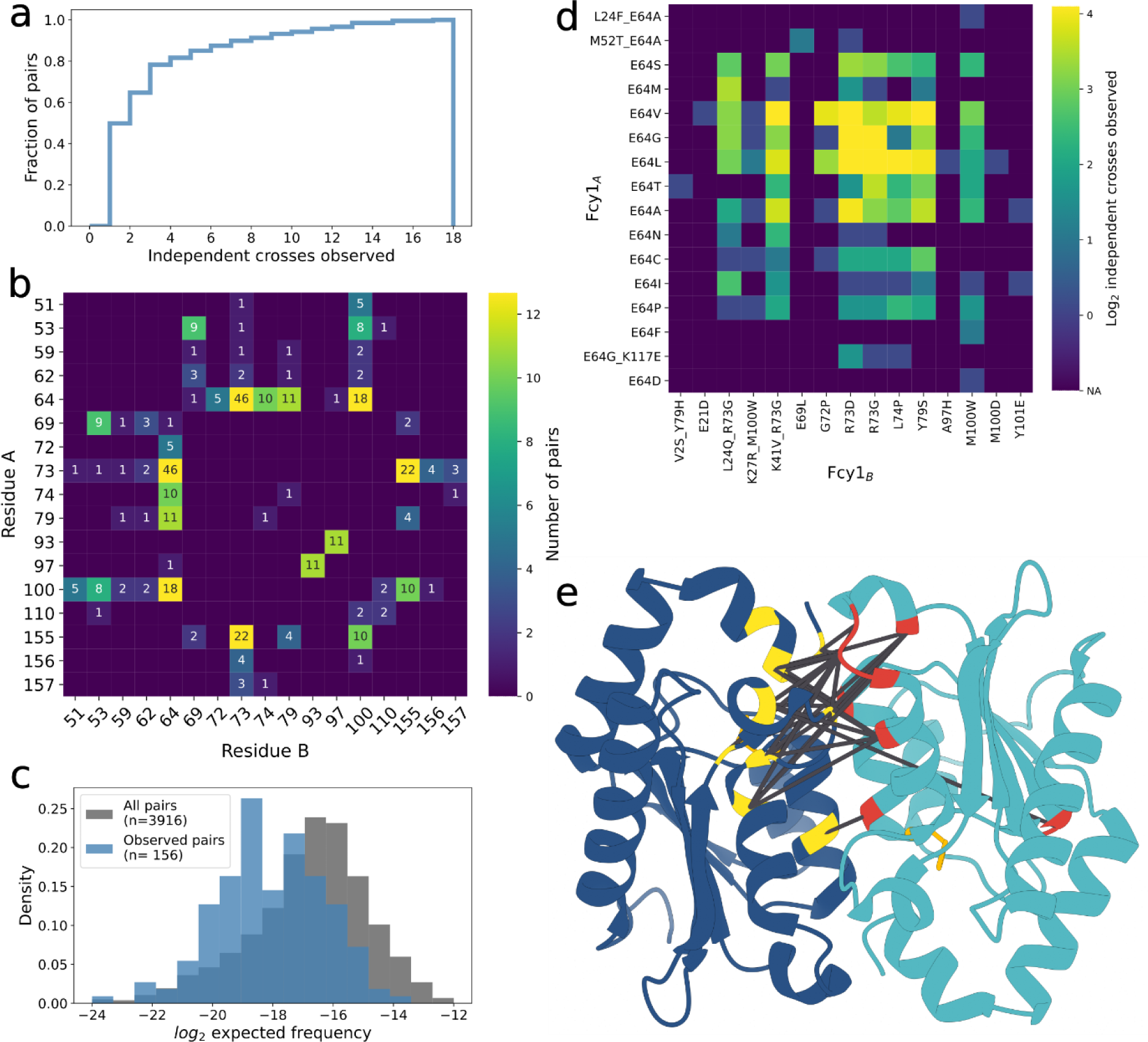
A network of positive *trans* epistatic mutations in a pair of duplicated gene copies. **a)** Cumulative distribution of independent observations for observed duplicate pairs with positive epistasis (n=207 pairs). **b)** Pairs of residues involved in at least two different *trans* interactions form a dense network of interactions. Color intensity denotes the number of observations for each pair with the numbers indicated each time. **c)** Predicted f requencies of a subset of Fcy1_A_-Fcy1_B_ pairs based on the initial abundance of Fcy1_A_ and Fcy1_B_ in the haploid LOF libraries. Of the subset of combinations involving any two *trans* interacting alleles we could predict, screen hits are shown in blue and unobserved mutant pairs are shown in grey. **d)** Positive *trans* epistatic network of E64 mutants. The substitutions observed are shown on the y axis with the mutants they associate with on the x axis. Color intensity denotes the number of observations for each pair. Double mutant alleles detected were also included when the causal mutation could be inferred (see methods). **e)** Hot spots of compensating mutation pairs (>2 occurences) mapped on the Fcy1 homodimer structure (PDB 1p6o). Residues proximal to their subunit’s active site are shown in yellow and those distal in red. Dark grey lines connect pairs of positions where positive epistatic mutations are found. The bound substrate analog and zinc cofactors are shown in orange.

When examining the distribution of compensatory mutation pairs in the Fcy1 structure, we found they were concentrated near the interface and active site, with 93% (192/207) occurring between variants among 16 positions (Figure 2b). This bias was not caused by differences in the initial abundance of variants and thus represents an enrichment due to the properties of these residues (Figure 2c, Figure S9a). Amino acid changes showing *trans* positive epistasis were generally also closer than expected to one another in the heterodimeric structure compared to random pairs and generally consisted of one interface mutant and one interface rim mutant (Figure 2e, Figure S9b-c). Surprisingly, we found that the majority (71%, 148/207) of pairs involved one variant whose LOF is likely caused by a loss of catalytic activity based on the known mechanism of reaction (*12–15*). For example, 13 non wild-type residues at the catalytic residue E64 were part of at least one compensating pair (Figure 2d). These most likely enzymatically dead mutants were frequently paired with the same set of other gene copies (e.g.: Fcy1_R73G_ or Fcy1_M100W_), suggesting a shared mechanism. A mutation at this site in one duplicate can therefore be broadly compatible with many other LOF alleles, increasing the probability of compatibility between mutants.

To validate these phenotypes, we reconstructed 18 gene sequences that were involved in 27 different complementing pairs in the initial experiment. We generated and assayed fitness individually all 324 possible duplicate pair combinations from these 18 variants. We recovered 90% (24/27) pairs from the initial experiment, and found three new pairs, supporting our analysis above suggesting that more epistatic combinations exist (Figure 3a).

**Figure 3:**
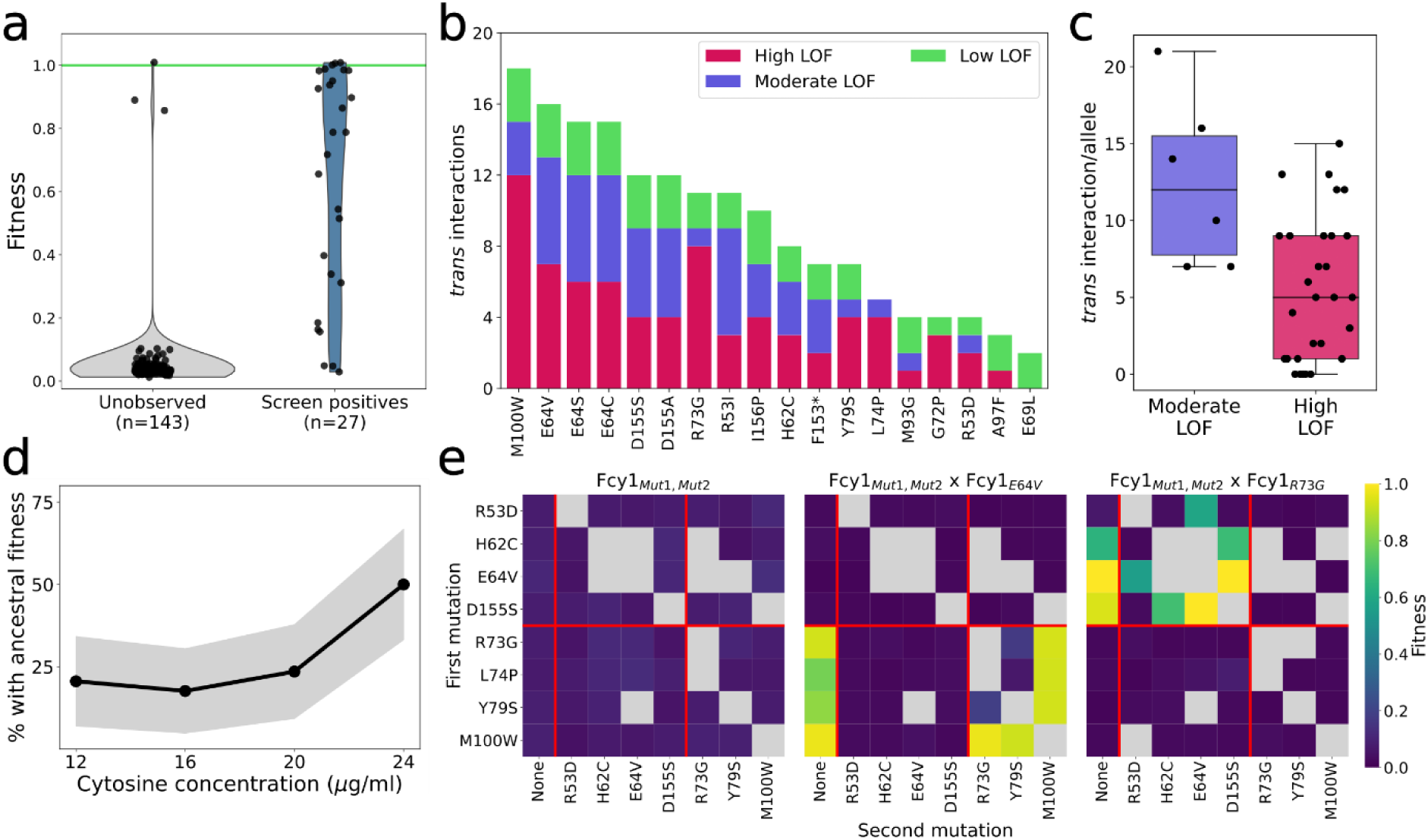
Compensatory mutations are frequent across the LOF landscape and often satisfy the requirements for constructive neutral evolution. **a)** Fitness (as measured by growth rate) of heterozygous mutants involving two distinct *FCY1* gene copies as a function of whether or not the combination of these sequences was detected as compensating pairs in the initial experiment. The green line represents the mean f itness of strains with a single wild -type gene copy. **b)** Number of compensatory mutant pairs observed for the 18 validation mutants, splitted between the different groups of LOF effect size. **c)** Frequency of *trans* interactions for moderate and high LOF mutations. Boxplots represent the upper and lower quartiles of the data, with the median shown as a grey bar. Whiskers extend to 1.5 times the interquartile range (Q3–Q1) at most. **d)** Proportion of pairs of gene duplicates (n=34 different pairs) with f itness comparable to the ancestral wild -type complex at different cytosine concentrations. The grey area shows the 95% confidence intervals around the proportion. **e)** Fitness of Fcy1 variants bearing two amino acid substitutions n *cis* (Fcy1_Mut1,Mut2_) alone or in combination with a *trans* interacting variant. Double mutant alleles were constructed sequentially, with the f irst mutations shown on the x axis and the second on the y axis. The red lines separate the control, the catalytic, and the interface mutations.

The results we present above include qualitative, nearly complete LOF mutations. However, more moderate LOF mutations could also be amenable to compensation, which would promote the maintenance of the heterodimers and extend the likelihood of the transition occuring by increasing the mutational target size. We therefore also examined if moderate LOF alleles could also participate in positive *trans* interactions between duplicates. To do this, we generated 20 other alleles with previously characterized functional effects, ranging from neutral variants to full LOF (*11*) and combined these pairs of gene copies together to test for compensation events (Figure S10-S12). We found new positive *trans* epistasis across the entire LOF gradient, both between and within different categories of effect sizes (Figure 3a and 3b). The network of positive *trans* effects is thus dense across the entire range of deleterious effects for Fcy1, vastly increasing the pool of mutations conducive to the transition to a heteromeric complex through compensatory changes.

A requirement for the non-adaptive replacement of a homomer by a heteromer following gene duplication is that the evolutionary transition to the heteromeric state should be neutral or nearly neutral. That is, the second compensatory mutation must restore activity enough to confer ancestral levels of fitness. To explore this, we compared the growth rates of 34 *trans* interacting allele pairs to the “ancestral state” single-copy gene. If our strains represent the post-duplication state, the equivalent ancestral state would be a haplosufficient diploid strain with a single gene copy (Figure S13). We found that ∼20% of gene pairs had growth indistinguishable from the ancestor (Figure 3d and S14, 6/34, p>0.1). The impact of a metabolic enzyme’s activity on fitness is likely to depend on environmental conditions, for instance substrate abundance. Some changes in environment conditions could therefore mask these slight defects in fitness and make the transition neutral. We tested this and found that increasing cytosine concentration by 50% was enough to increase the fraction of gene pairs with wild-type-like fitness to half of the newly evolved heteromers tested (17/34). Our results show that compensation between LOF mutations can occur neutrally or nearly neutrally in regards to the ancestral level of function.

Our evolutionary model supposes that the second LOF mutation occurs in the second gene copy, leading to the restoration of function through the production of a heterodimeric complex. However, the second mutation would have the same probability of occurring in the first gene copy. If positive epistasis could also occur in *cis* for these same mutations, this second mutation could also restore function and favor the maintenance of two functional homodimers and eventually the return to a single copy. We therefore examined if the epistatic interactions we uncovered were specifically acting in *trans*. We constructed a set of 23 *cis* double mutant genes combining LOF mutations that positively interact in *trans* (Figure 3e). None of the *cis* double mutants restored growth, independent of whether or not the same combination showed a positive *trans* interaction. Interestingly, some of these double mutant *FCY1* copy maintained the ability to compensate another single mutant copy in *trans*. This suggests that two strongly deleterious mutations within one gene copy can occur without preventing the emergence of downstream positive *trans* epistatic mutations.This could greatly increase the evolutionary window where constructive neutral evolution can occur.

Having shown that many *trans* epistatic interactions between Fcy1 LOF mutants allow the functional replacement of a homodimer by a heterodimer, we investigated the molecular mechanisms. We verified three assumptions that are critical for our model. First, that the effects are not dependent on both gene copies being expressed at the *FCY1* locus, for instance through regulatory feedback loops, as a duplication would have placed one copy at the non-homologous position. To test this, we engineered a haploid strain with two copies of *FCY1*, one at the endogenous locus and the other under the control of an inducible promoter (figure S15a). This allowed us to validate both that the genomic context of the *FCY1* locus is not necessary for *trans* epistasis and that compensation is expression dependent (Figure S15b-c).

Second, we verified that the two proteins produced by duplicated pairs that restore the ancestral function indeed form heterodimers in living cells. For this, we used an assay that reports on complex formation in living cells, the DHFR-PCA protein fragment complementation assay (*16*). We detected all (6/6) expected heteromeric complexes involving Fcy1_E64V_ or Fcy1_R73G_ (Figure S16), showing that mutant subunits indeed assemble into complexes in cells. Interestingly, Fcy1_R73G_ could form stable dimers with Fcy1_E64V_ but not with itself nor the wild type, while Fcy1_E64V_ interacted with both. This result suggests Fcy1_E64V_ might actually act to stabilize Fcy1_R73G_, revealing a potential mechanism of compensation.

Our third assumption was that the heteromers themselves have enzymatic activities and that these activities are not dependent on some feedback mechanisms or moonlighting activities of other enzymes that would take place in yeast cells specifically expressing LOF gene copies. To test this, we purified individual Fcy1 variants and their homomer-heteromer mixtures by co-expressing them together in bacteria. While none of the 5 LOF mutants tested had detectable activity when expressed alone, all pairs of gene copies that compensated each other in yeast were active (Figure 4a). The activity of these homomer- heteromer mixtures is lower than the wild-type enzyme expressed alone but seems to result in comparable growth rates *in vivo*. One possible explanation is that depending on the limiting step of the metabolic pathway involved, lower activity might not result in reduced growth rate if cytosine deamination is not usually the control step (*7*). Also, it is important to note here that in these mixtures, only a fraction of the complexes are expected to be heteromers: 50% if both the homomeric and the heteromeric complexes are equally stable and both proteins have the same abundance (*5*). Any mechanism that could specifically stabilize the heterodimer or degrade an unstable subunit *in vivo* could modulate this ratio. For instance, chaperones that could buffer part of the loss of stability would increase the amount of functional protein complex in the cell and help bring fitness to that observed in the wild-type.

**Figure 4:**
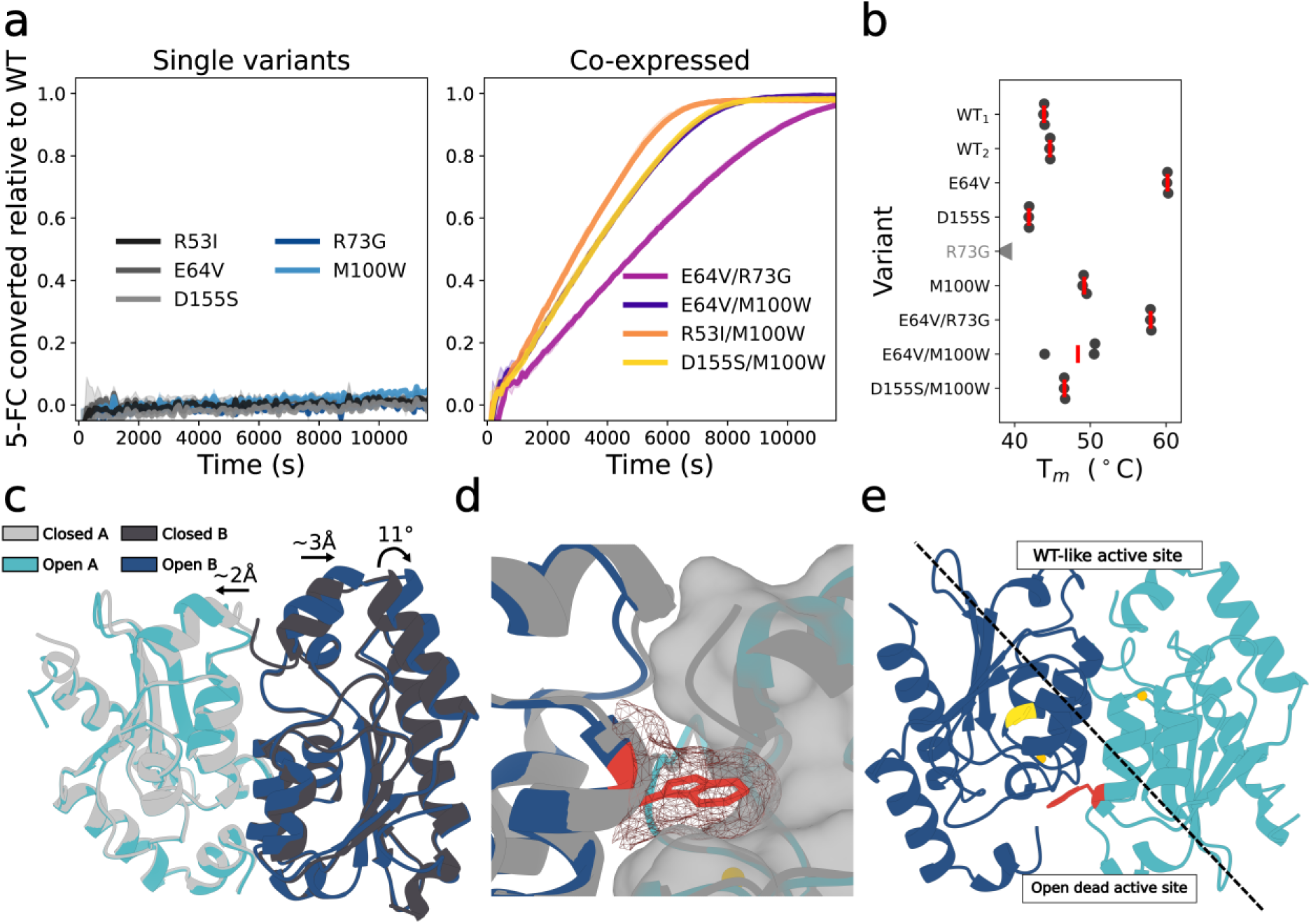
Newly active heteromers are active *in vitro* and show signs of subfunctionalization. **a)** Relative activity of homomer-heteromer mixtures as well as the individual variants. The curves show the mean of three technical replicates, with the shaded area representing the 95% confidence interval. **b)** Stability of LOF variants alone and in combination and homomer-heteromer mixtures as measured by nanoDSF. The red line indicates the median of three technical replicates. Fcy1 _R73G_ is shown in grey as it was already unfolded by the start of the assay, and so is assumed to have a lower T_m_ than the wild-type. **c)** Superposition of the open and closed structures of the Fcy1 dimer (open: PDB 8VLK, closed: PDB 8VLJ), showing the movement of the G72-K80 loop and the C-terminal helix on one end of the dimer. **d)** Steric clash between the W100 side chain and the C-terminal helix (closed form) of the other subunit. The tryptophan sidechain prevents the C-terminal helix f rom adopting the closed conformation required for catalysis. **e)** Mechanistic model for the compensatory relationship between Fcy1_E64V_ and Fcy1_M100W_. While the W100 sidechain (red) prevents the C-terminal loop of the E64V(yellow) subunit to close on the active site, this catalytic site is already inactive and so is not affected by this additional disruption. On the opposite side of the dimer, a wild -type like catalytic site can form due to both deleterious mutations being concentrated in space.

Having validated that the positive *trans* epistastic effects we observed are due to heteromerization in vivo that leads to direct molecular compensation, we investigated the properties of variants forming active complexes. We measured the stability of both homomer and homomer-heteromer mixtures (Figure 4b). As suggested by our experiments validating the assembly of heteromeric complexes *in vivo* (Figure S16), the catalytic dead mutant Fcy1_E64V_ showed a dramatic increase in T_m_ compared to the wild-type (p=5.4x10^-10^, two-sided Welch’s t-test). As expected, Fcy1_R73G_ was less stable than the wild-type as it was already denatured at room temperature when we started measurements. The other presumed catalytic dead mutant, Fcy1_D155S_, was less stable than the wild-type (p=3.6x10^-5^, two-sided Welch’s t-test), suggesting that increased stability of the catalytically inactive subunit is not a prerequisite for heteromer assembly in all cases. Similarly, Fcy1_M100W_ was more stable than the wild-type (p=7.43x10^-7^, two-sided Welch’s t-test), showing that LOF independent of catalytic residues is not necessarily associated with losses in stability.

To try to rationalize *trans* epistatic effects, we examined the Fcy1 homodimeric enzyme structure. The loop from G72 to K80 at the dimer interface is enriched in *trans* interacting residues (G72, R73, L74, Y79). This interface loop is in close contact with the C-terminal alpha-helix of the other subunit, which is predicted to move as part of substrate entry and subsequent product release (*12*, *14*, *15*). Importantly, the C-terminal helix also contains residue D155, which directly stabilizes both the substrate and the H62 sidechain in the conformation required for catalysis (*17*). Mutations that disrupt the loop helix interactions could thus disrupt the open to closed state transition and substrate positioning for catalysis, explaining the LOF phenotype of loop mutant homomeric complexes. To validate the importance of this conformational change, we solved the first crystal structure of Fcy1 in its open form (Figure 4c, table S1). This new structure shows the terminal C-helix indeed detaches from the dimer interface in the open conformation, with the last two residues becoming disordered. Homomeric complexes of loop mutants are thus unable to complete the catalytic cycle, as well as sometimes being unstable.

Importantly, we also observed that only one subunit at a time appears to adopt the open conformation, with the other staying closed. The lack of necessity for coordinated movement between the two active sites of the dimer could act to insulate them from one another in *trans* interacting heteromers. This would result in a wild-type like active site on one end of the complex while the other is inactive due to the catalytic residue mutation (Figure 4d).

Because most of the loop mutants (like Fcy1_R73G_) were too unstable for *in vitro* characterization, we focused on another frequent and stable *trans* interacting mutant, Fcy1_M100W_. We hypothesized that for this mutant, the change to a bulky tryptophan sidechain could physically prevent the C-terminal helix from returning to the closed state and thus disrupt catalysis. The crystal structure of the Fcy1_M100W_/Fcy1_M100W_ homodimer (Figure 4d) shows this is indeed the case, with both active sites stuck in the open state. In this “permanently open” structure, the important residue D155 is displaced by 2.3-2.5 Å from its catalytically competent conformation and, as a result, can not properly orientate the substrate for catalysis. Finally, we solved the structure of the Fcy1_E64V_/Fcy1_M100W_ heterodimer (Figure 4e). The structure of the heterodimer shows that the active site harbouring the M100W and E64V mutations is indeed locked in the open conformation while the other insulated active site adopts a wild-type like closed conformation compatible with the dynamic conformational switch required for a complete catalytic cycle. Through the separation of Fcy1 active sites in space and of their catalytic cycle in time, LOF variants can readily assemble into active heterodimers. This effect can be compounded by changes in stability of the different subunits, setting the stage for further specialization downstream.

Our results show that strong *trans* epistasis between mutations that result in loss of activity in nascent paralogs can be both frequent and of large magnitude, providing numerous avenues for the replacement of obligate homodimers by obligate heterodimers. The fact that very few molecular changes are needed without the need for positive selection makes it possible for this mechanism to occur in nature. Because gene duplication occurs at high frequency (*18*) and many gene duplications in themselves have no deleterious effects (*19*), duplicates will often segregate in large populations and provide the raw material for this transition to happen. How frequently this transition effectively occurs will remain to be examined but some detailed examples are consistent with this model. For example, ancestral sequence reconstruction of the fungal V-ATPase inner membrane ring suggests that similar events could have led to an increase in the number of subunits required for complex assembly (*8*). Our results also illustrate how two paralogous copies can quickly become co-dependent following the accumulation of positive *trans* epistatic interactions following duplication. Recent studies on protein-protein and genetic interaction network analysis have shown that many paralog pairs have become co-dependent to accomplish a single function (*9*, *20*, *21*). We show that only one mutation per paralog is sufficient to induce such physical and functional co-dependency. Over longer evolutionary timescales, each gene copy could specialise further, resulting in paralogous subunits with well defined functions. This has been observed for instance in pairs of kinases whereby one of the two paralogs has become a regulatory subunit of the other after having lost its catalytic activity (*22*). Such constructive neutral evolution could therefore sometimes serve as a launchpad for protein sequence space exploration leading to new molecular functions.

## Methods

### Reagents and strains

Media recipes are detailed in table S2. Unless stated otherwise, all yeast cultures were incubated at 30°C. Oligonucleotides, plasmids and yeast strains used are presented as tables S3, S4 and S5 respectively and are available upon request.. Cytosine and 5-FC were purchased from Alfa Aesar (now Thermo Scientific Chemicals, Haverhill, MA, USA), lot number B4046A and L16496 respectively.

### Systematic mutagenesis library generation

We used the same approach as in Després et al, 2022 (*11*) to generate Fcy1 variant libraries in a MATa and a MATα strain in parallel. In this study, we created mutant plasmid libraries for each *FCY1* codon covering all possible alternative codons. We then amplified this pool of *FCY1* alleles to serve as donor DNA for genome editing with CRISPR-Cas9 to insert mutants at the endogenous *FCY1* locus. To facilitate diploid selection, we generated new receiver strains with different selection markers for each mating type. Starting from strains where *FCY1* was deleted with an antibiotic cassette (BY4741 *fcy1Δ::NatNT2* and BY4742 *fcy1Δ::HphNT1*, respectively s_001 and s_002), we replaced the antibiotic cassette with a short sacrificial sequence (stuffer3) as in Dionne et al (*23*) using a CRISPR-Cas9 transformation (*24*) and cassette specific pCAS vectors (p_001 and p_002) to create markerless deletion strains (BY4741 *fcy1Δ::stuffer3* and BY4742 *fcy1Δ::stuffer3*). We then replaced the HO locus of both strains with an antibiotic marker to allow for easier diploid selection, resulting in our final receiver strains s_003 (BY4741 *fcy1Δ::stuffer3 hoΔ::NatNT2)* and s_004 (BY4742 *fcy1Δ::stuffer3 hoΔ::HphNT1*).

For each codon, we then amplified the corresponding mutagenized allele pool from our plasmid libraries using the following PCR reaction:

| Reagent | Volume (μL) |
| --- | --- |
| Kapa HF buffer 5X | 5 |
| dNTPs 10 mM | 0.75 |
| Oligo knock_in_for 10 μM | 0.75 |
| Oligo knock_in_rev 10 μM | 0.75 |
| Kapa polymerase | 0.17 |
| Variant plasmid library 2 ng/μL | 5 |
| PCR grade water | 12.58 |
| Total | 25 |

With the following PCR cycle:

|  |  |  |
| --- | --- | --- |
| 95 °C | 5 min |  |
| 98 °C | 20 s | 29X |
| 60 °C | 15 s |  |
| 72 °C | 45 s |  |
| 72 °C | 1.5 min |  |

We then performed one transformation/codon/strain using the same modified CRISPR-Cas9 transformation protocol for 96 well plates we used to construct the previous Fcy1 mutagenesis library (*11*, *23*, *25*). Each well contained 20 μL competent cells of the appropriate strain, 2 μL carrier DNA (salmon testes ssDNA 10 mg/mL, boiled for 5 min and then cooled on ice) and 3.75 μL of 200 ng/μL pCAS-Stuffer3, prepared as a master mix and dispensed before adding 10 μL donor DNA. We then dispensed 200 μL of freshly prepared PLATE solution (40% PEG 3350, 100 mM lithium acetate, 1 mM Tris-Cl, 0.1 mM EDTA, filter sterilized) in each well. The plates were sealed with sterile aluminium foil, vortex mixed and incubated at 30 °C for 30 min before a 15-min heat shock at 42 °C. The plates were then centrifuged at 448g for 6 min to pellet the cells, the supernatant was removed, 100 μL YPD was added and the plates were sealed with aluminum foil again for 3.5 h outgrowth at 30 °C. The cells were then plated on YPD+G418+NAT or YPD+G418+Hyg depending on the strain and incubated at 30 °C for ∼64 h.

We recovered cells from each transformation by soaking the plates with 5 mL YPD before scraping them with a glass rake. The cell suspension was then transferred to a 15 mL falcon tube (one per mutated residue), vortexed and the OD was measured. Codon-specific transformations were combined in three libraries covering overlap regions of the coding sequence as previously by mixing 5 OD units of each transformation (pool 1 contains codons 2 to 67, pool 2 codons 49 to 110 and pool 3 codons 93 to 158). The final OD of each pool was measured to get cell concentration, and multiple glycerol stocks per pool were prepared (950 μL cells mixed with 450 μL 80% glycerol) and stored at -80 °C until use. Each codon-specific transformation was also kept as a glycerol stock in a 96-well plate (same recipe, adjusted to 250 μL final volume). Cell pellets were also kept for downstream DNA extraction and sequencing.

### Selection of loss-of-function mutants

For each mating type, the three replicate *FCY1* fragments pools were thawed on ice and used to inoculate 5 mL of SC complete media in triplicate with 2.5 OD units (>5,000X coverage of library mutants) and grown overnight at 30°C with shaking. The following morning, each pool was diluted to 0.15 OD in 5 mL (∼1,750X library coverage) SC + 100 μg/mL 5-FC and grown for 24h, so that ∼6 generations had elapsed when the culture reached saturation. The pools were passaged a second time in the same conditions and then stored as glycerol stock. To test selection efficiency for LOF mutants, we inoculated 3 mL SC -ura + 50 μg/mL cytosine cultures 0.1 OD from each pool. We observed growth in a large fraction of test cultures, and so decided to perform two other rounds of 5-FC selection to ensure efficient purging of functional *FCY1* alleles. The cells from the second passage were thawed on ice and inoculated at 0.3 OD in SC complete media. The next morning, cells were diluted to 0.15 OD in 5 mL SC + 100 μg/mL 5-FC for two other passages, leading to the final count of ∼24 generations under strong negative selection. The final OD of each pool was measured to measure cell concentration, and multiple glycerol stocks per pool were prepared (950 μL cells mixed with 450 μL 80% glycerol) and stored at -80 °C until their use. Cell pellets were also kept for downstream DNA extraction and sequencing.

### Variant library sequencing

To validate our variant libraries and to measure allele frequency changes after negative selection, we used the same approach developed for the first systematic mutagenesis screen of Fcy1 (*11*). DNA was extracted from each pool’s cell pellet using a standard phenol/chloroform method (*26*). The appropriate region of *FCY1* was then amplified by PCR with fragment-specific oligonucleotides with the following reaction:

| Reagent | Volume (µL) |
| --- | --- |
| Kapa HF buffer 5X | 3 |
| dNTPs 10 mM | 0.45 |
| Pool forward oligo 10 µM | 0.45 |
| Pool forward oligo 10 µM | 0.45 |
| Kapa polymerase | 0.1 |
| Variant pool gDNA 30 ng/µL | 5 |
| PCR grade water | 5.55 |
| Total | 15 |

With the following PCR cycle:

|  |  |  |
| --- | --- | --- |
| 98 °C | 3 min |  |
| 98 °C | 30 s | 20X |
| 60 °C | 15 s |  |
| 72 °C | 30 s |  |
| 72 °C | 1 min |  |

After gel migration, the amplicons were diluted 1/2,500 by two consecutive 2 μL PCR: 98μL PCR grade water dilutions. This was then used as a template for a second PCR round where sample-specific barcodes were added at both ends of the amplicon using the Row-Column PCR approach (*27*), with each reaction performed in duplicate.

| Reagent | Volume (μL) |
| --- | --- |
| Kapa HF buffer 5X | 3 |
| dNTPs 10 mM | 0.45 |
| Row-Forward oligo 2 μM | 2.25 |
| Col-Reverse oligo 10 μM | 0.45 |
| Kapa polymerase | 0.1 |
| Diluted PCR | 2.25 |
| PCR grade water | 6.5 |
| Total | 15 |

With the following PCR cycle:

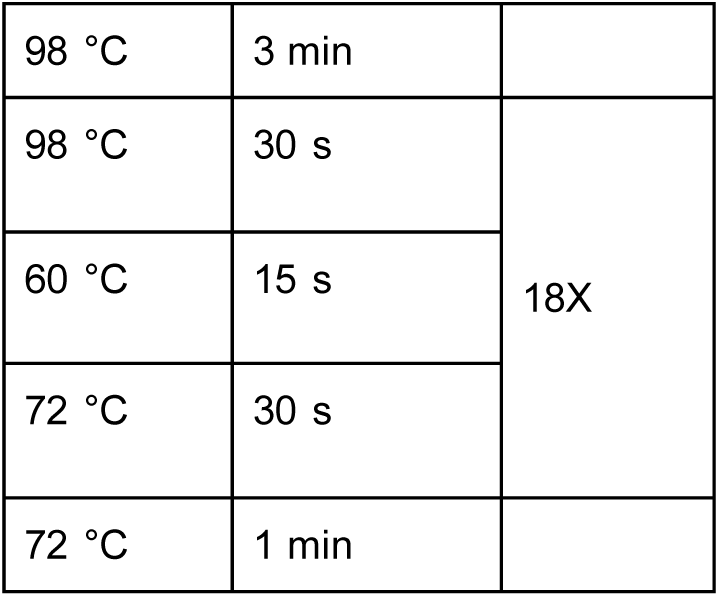

Replicate reactions were pooled together, purified using magnetic beads, and quantified using a NanoDrop spectrophotometer. Barcoded amplicons of the same pools were then mixed in equivalent quantities. The resulting pools were diluted and used as templates for a final PCR round performed in quadruplicate to add plate barcodes at each end as well as the Illumina p5 and p7 sequences:

| Reagent | Volume (μL) |
| --- | --- |
| Kapa HF buffer 5X | 3 |
| dNTPs 10 mM | 0.45 |
| Plate_for-Forward oligo 10 μM | 0.45 |
| Plate_rev-Reverse oligo 10 μM | 0.45 |
| Kapa polymerase | 0.1 |
| Purified RC-PCR pool 0.01 ng/μL | 1 |
| PCR grade water | 9.55 |
| Total | 15 |

With the following PCR cycle:

|  |  |  |
| --- | --- | --- |
| 98 °C | 3 min |  |
| 98 °C | 30 s | 20X |
| 61 °C | 15 s |  |
| 72 °C | 35 s |  |
| 72 °C | 1 min |  |

The replicate reactions were pooled, purified on magnetic beads and gel verified to obtain the final libraries. We performed two rounds of sequencing: the first to verify sequence diversity in the variant library pools, and the second to measure allele frequencies after negative selection. Both sets of samples were sequenced using the MiSeq Reagent Kit v3 on an Illumina MiSeq for 600 cycles (IBIS sequencing platform, Université Laval).

Variant abundance and log_2_ fold-changes were computed using the same approach as in Després et al. (*11*). We followed the standard practice of adding one to all variant counts to make fold-change calculations possible for variants that dropped out during the competition. Variant read counts were then divided by total read count (including the wild-type sequence) to obtain relative abundances. Relative abundances were then compared between the end of the SC media passage and the end of the 5-FC passages to calculate log2 fold-changes for all mutant codons. We discarded all codon-level variants that were covered by less than 20 reads before selection. To convert to amino acid log2 fold-change, we considered each group of synonymous codons as replicates and calculated the median fold-change of each group. We then scaled log2 fold-changes to the difference between nonsense and synonymous codons so that:

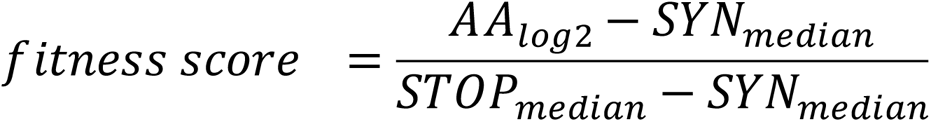

We took the median score of each Fcy1 variant across library replicates in the same mating types and calculated the final fitness score by taking the median of all such scores for each variant (the overall median). Finally, we also calculated a score based only on data from the variant pools that would be used in the mating step. We used the distribution of scores for synonymous mutations to set the threshold at which we classified an amino acid variant as nonfunctional (Figure S5d). This cutoff resulted in the identification of 1071 LOF missense variants. Fitness scores of variants are provided as table S6.

### Mating libraries and pooled auxotrophy complementation assay

Crossing all strains expressing *FCY1* fragments against the other results in a 3x3 matrix where the off-diagonal crosses are replicated twice (e.g. Pool 1 MATa x Pool 2 MATα is a biological replicate of Pool 2 MATa x Pool 1 MATα). To have each cross in duplicate, we performed another set of crosses on the diagonal using a different 5-FC passaging replicate for the Matα LOF pool. We also performed two pairs of control crosses against either a *FCY1* deletion strain (negative control) or a wild-type *FCY1* allele (positive control). We used the same mating and diploid selection protocol used previously for protein-protein interaction assays of variant libraries (*25*).

The pools of LOF mutants were thawed on ice and ∼1 OD unit (>5,000x coverage of LOF variants diversity) was used to inoculate 20 mL pH buffered SC (MSG) overnight cultures with antibiotic selection (100 μg/mL nourseothricine for MATa libraries or 250 μg/mL hygromycin B for MATα libraries). The saturated cultures were spun down and resuspended in water to a concentration of 25 OD/mL. We then mated mutant pools together by inoculating 50 ml YPD cultures in flasks with 25 OD of each parental library, resulting in 74x- 160x coverage of LOF *FCY1* variant pairs. After 8 hours of mating, the cells were spun down, and resuspended in 5 mL sterile water and 25 OD units were transferred to 50 mL YPD + 100 μg/mL nourseothricine + 250 μg/mL hygromycin B flasks for a first diploid selection round. After 24 hours, cells were spun down and resuspended in 5mL sterile water, and 25 OD units of cells were transferred to SC (MSG) + 100 μg/mL nourseothricine + 250 μg/mL hygromycin B for the second round of diploid selection. After 18h, cells were spun down and resuspended in water. For each cross, we then inoculated 100 mL SC-Ura + 16 μg/mL cytosine with 25 OD units of diploid pools, resulting in a final cell density of 0.25 OD/mL. The remaining cells were mixed with glycerol 80% and stored at -80°C.

After 18h of enrichment for Fcy1 function in cytosine media, optical densities for LOF x LOF and LOF x *fcy1Δ* crosses were close to the baseline measurements, as expected if the vast majority of LOF mutant pairs do not complement the auxotrophy, while the WT x LOF crosses had already saturated. For each cross, we plated 100 ul cell culture and 100ul of a 1/10 dilution in duplicate for each cross on SC-Ura + 16 ug/mL cytosine. After 48h incubation, pictures of the plates were taken and CFU/mL was estimated with the OpenCFU colony counting software (*28*) using default parameters. We picked 96 colonies per plating (2 plates/cross, 24 total) and patched them on SC complete media to array them in a multichannel compatible configuration. After 48h, cells from patches were resuspended in 200 μL sterile water in Greiner flat bottom 96 well plates. The cell suspension was transferred to 96-well PCR plates, spun down, and kept at -80°C for downstream DNA extraction.

### High throughput single colony *FCY1* genotyping

To extract DNA for *FCY1* genotyping of the putative heterozygous compensating strains, we used a modified version of the Quick DNA method (*29*). In each well of the PCR plates, we resuspended cell pellets in 100 μL 200 mM LiOAc, 1% SDS, sealed the plate and incubated it for 15 minutes at 70°C in a thermocycler. We then added 150 μL ice-cold 95% ethanol and centrifuged the plates at 2,204g for 20 minutes. The supernatant in the plate was then removed, and the pellet was washed with 150 μL ice-cold 70% ethanol and then spun again at 2,204g for 20 minutes. The supernatant was removed and the pellet was dried in a dry stove set at 55°C. The dry pellet was then resuspended in molecular grade water to resolubilize the DNA and cell debris was spun down 1 min 20 s at 2,204*g*.

To amplify the *FCY1* locus, we used two pairs of oligonucleotides amplifying the entire coding region while also adding flanking sequence with homology to Illumina Nextera adapters. To reduce the number of index pairs required to barcode individually the 2,304 mutants we isolated, we used slightly different oligonucleotides for even and odd plates. The second PCR oligonucleotides for the even plates added 3 degenerate bases between the *FCY1* primer binding site and the start of the Nextera adapters so that amplicons with the same index pairs could be distinguished after sequencing (Nextera_i5-FCY1-Nextera_i7 vs Nextera_i5-3N-FCY1-3N-Nextera_i7). All PCR steps until pooling were performed using filtered tips to avoid cross-contamination. We amplified the *FCY1* alleles from the extracted DNA using the following reaction:

| Reagent | Volume (μL) |
| --- | --- |
| BioShop TAQ buffer 5X | 4 |
| dNTPs 25 mM | 0.16 |
| 0N/3N forward oligo 10 μM | 0.4 |
| 0N/3N reverse oligo 10 μM | 0.45 |
| BioShop TAQ | 0.24 |
| DNA (Quick DNA extraction) | 2 |
| PCR grade water | 12.8 |
| Total | 20 |

With the following PCR cycle:

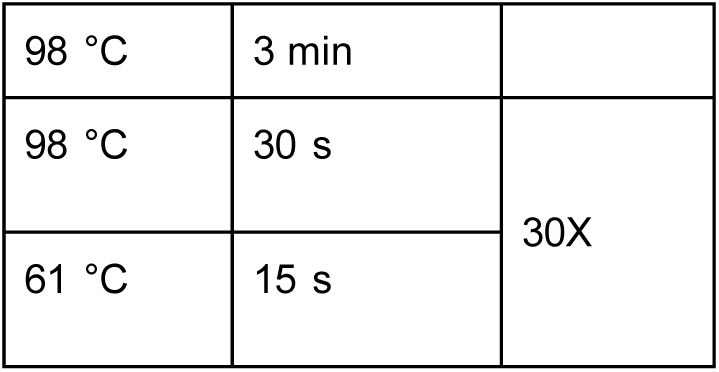
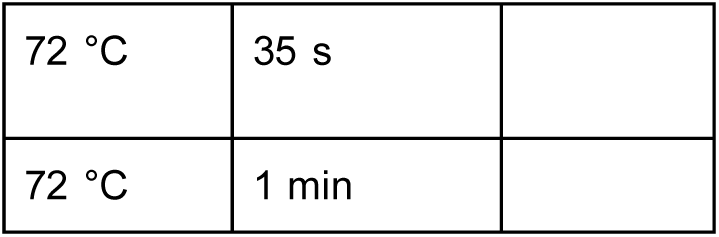

The PCRs products were stored at -20°C until use for the second PCR. After thawing, the products were diluted 1/2,500 through successive 2:98 dilutions and immediately added to the indexing PCR mix:

| Reagent | Volume (µL) |
| --- | --- |
| Kapa HF buffer 5X | 3 |
| dNTPs 10 mM | 0.45 |
| Nextera_i5_BC_For oligo 5 µM | 0.90 |
| Nextera_i7_BC_Rev oligo 10 µM | 0.45 |
| Kapa polymerase | 0.1 |
| Diluted product | 2 |
| PCR grade water | 8.1 |
| Total | 15 |

With the following PCR cycle:

|  |  |  |
| --- | --- | --- |
| 98 °C | 3 min |  |
| 98 °C | 30 s | 18X |
| 61 °C | 15 s |  |
| 72 °C | 30s |  |
| 72 °C | 1 min |  |

Indexed amplicons from each plate were pooled together and purified using magnetic beads. The DNA concentration of each plate pool was measured using a NanoDrop spectrophotometer and used to adjust the amount of each plate library in the master pool sent for sequencing. The libraries were sequenced using the MiSeq Reagent Kit v3 on an Illumina MiSeq for 600 cycles (CHUL sequencing platform, Université Laval)

### Detection and filtering of complementing mutant pairs

The sequencing libraries were processed using custom Python scripts. Briefly, forward and reverse reads pairs were merged using panda-seq (*30*). The merged reads were then filtered by total read length using vsearch (*31*) to only keep reads matching the length of the *FCY1* coding sequence, taking into account the presence or absence of the 3N sequence at the end of the amplicons. This resulted in the final set of 2,304 demultiplexed sequences corresponding to each isolated colony. We then clustered the reads within each sequence file to 100% identity using vsearch, allowing us to obtain counts for all unique sequences within each file. We detected mutations by directly comparing the sequenced alleles with the wild-type codon-optimized *FCY1* sequence. We then applied a series of filters to reduce noise introduced by library preparation failures or false positives linked to leftover functional *FCY1* alleles that might still be present after selection:

1. We excluded all samples for which we could not recover over 250 reads.
2. We filtered out all samples for which the top 2 alleles represented less than 10% of the total reads.
3. We filtered out samples where one of the two most abundant alleles was far more abundant than the other by setting a threshold on the abundance ratio of the top two alleles so that reads_allele 2_ / reads_allele 1_ > 0.4
4. We filtered out any sample where the total number of mutations across both alleles is more than 4
5. We removed any sample where one of the top alleles is either the wild-type or a mutation with a fitness score ≤ 0.31 (the 97th percentile of synonymous mutants).

Additionally, we excluded 6 pairs containing one of two double mutant alleles (Y79I_K80M and G76V_G118D) because the fitness scores scores of both K80M and G118D were inconsistent between the two 5-FC systematic mutagenesis assays (fitness score_100_ > fitness score_1.56_).

This stringent filtering resulted in a set of 887 high-confidence LOF x LOF positive *trans* epistatic events, spread across 207 unique Fcy1 variant pairs (table S7). Rarefaction curves were generated by taking the median of 100 iterations per curve. To obtain the number of independent crosses for each variant pair, we summed the number of unique codon combinations in each mating pool resulting in the same amino acid combinations. To calculate the predicted abundance of each heterozygous strain in a given cross, we used the following equation:

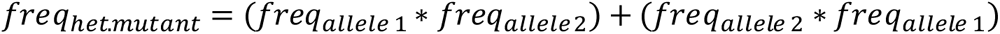

We then summed these values across all crosses. All variant pairs identified were observed in the expected cross based on the region mutagenized in their parental pools, although some double mutant alleles we detected included amino acid changes outside this region (ex.: Fcy1_K41V+R73G_, present in 12 pairs). To generate random distributions of LOF allele pairs, we drew 207 random pairs of mutants with fitness score_100_ > 0.31, sampled from each mating pool in the same proportion as what was observed in the high-confidence heterozygous mutants set. We performed this over 1,000 iterations to generate a null distribution of expected properties in LOF allele pairs. This was repeated for each variant property of interest. The full list of unique allele pairs is provided in table S7.

### Fcy1 variant effect prediction and structure analysis

We used the same in silico methods to calculate the effects on structure and interface stability of Fcy1 mutants as previously (*11*). Briefly, we used the crystal structure from Ireton et al ((*12*), PDB 1p6o) to compute residue relative solvent accessibility (RSA), distance to the active site and interface, and distance between residues within and between chains. The distance to the active site was considered to be the shortest distance between a specific amino acid and any active site residue identified in (*12*). Then, the FoldX BuildModel and AnalyseComplex functions were used to estimate mutational effects on ΔG. RSA and temperature measurements of both chains were averaged to obtain the final value. To determine which residues were part of the interface, we used a definition based on the distance to amino acids in the other subunit, where residues whose two closest non-hydrogen atoms are separated by a distance smaller than the sum of their van der Waals radii plus 0.5 Å are considered to be at the interface (coded as 1). Residues whose alpha carbons are within 6 Å of the other chain were considered to be at the interface rim (near the interface, coded as 0.75).

### Validation strain construction

We selected 18 alleles observed in our dataset, as well as 30 other *FCY1* variants we had previously characterized as part of validation experiments for our previous assays (*11*). We used the same CRISPR-Cas9 knock-in strategy as in the variant library generation, starting from the same haploid deletion strains s_005 (MATa) and s_006 (MATα). We generated mutant donor DNA using a fusion PCR strategy where oligonucleotides containing the mutations of interest are used to generate two overlapping amplicons bearing the mutation that are then fused into a second PCR. The first reaction, which was amplified from the five-prime flanking homology arm before the start of the coding sequence up to the mutated amplicon overlap, used oligonucleotides fusion_for and mutant reverse. The second reaction, which amplified from the mutated overlap to the three-prime homology arm after the coding sequence, used oligonucleotides mutant forward and fusion reverse. Both resulting amplicons were then diluted 1:2,500 and used as the template for the second reaction using oligonucleotides fusion_for and fusion_rev to reconstitute the full-length *FCY1* sequence. We used the same approach to later construct double mutants, but using genomic DNA of single mutants as the starting template instead of the wild-type gene.

We then performed one-by-one knock-ins using the same reaction setup as for the variant library, but using the fusion PCR products as donor DNA. Transformants (four per mutant) were used to inoculate 1.2 mL YPD cultures, grown overnight, and then streaked on YPD to purge the pCAS plasmid. One colony from each streak was resuspended in 200 μL water and spotted on YPD as well as YPD + G418. *FCY1* integration was verified by PCR using oligonucleotides FCY1_alt_A and fusion_rev on quick DNA extractions and PCR-positive strains were then validated by Sanger sequencing using fusion_for as the sequencing oligonucleotide. We successfully constructed 41/48 MATa and 46/48 MATα strains: these were then stored as glycerol stocks until used. To validate the fitness score values and validate loss of growth in cytosine media for all variants, we performed growth curves in SC + 100 ug/mL 5-FC, SC -Ura + 200 ug/mL cytosine and SC -Ura + 20 ug/mL cytosine. We also included a wild-type strain and both deletion strains as controls. Cells from overnight SC media precultures were spun down and resuspended in water at 1 OD. We then inoculated 180 ul of 1.11X media + 5-FC or cytosine with 20 ul cells, bringing the final cell density to 0.1 OD. Measurements were taken every 15 minutes over 30 hours using a Tecan Infinite M Nano (Tecan Life Sciences) at 30°C. The *MATa* strain bearing the H62C mutation was excluded from further analysis as the growth curves showed it to be a mix of two genotypes.

### Mating and growth rate measurements

To perform all against all mating and growth curves, the validated strains were revived from the stock plates on YPD+Nat or YPD+Hyg as appropriate in 6 copies using an S&P BM3-SC robot (S&P Robotics). We randomized the position of each one of the 2,016 (42*48) matings by assigning a position to each mating on one of six 384 colony arrays (quadrants C and D of plate 6 were empty). We rearrayed the haploid strains on the appropriate media using the cherry-picking tool. We then mated the cells by printing both parental arrays on the same YPD plate and incubated for 48h. Diploids were selected by replicating the mating arrays on YPD+Nat+Hyg twice. To perform the growth measurements, we used the 96-pin replication tool to deposit cells from a 384 array quadrant in 96-well flat bottom plates (Greiner) filled with sterile water. The cells were resuspended using a 96-well plate shaker, and OD was measured: cell concentrations were between 0.4-0.7 OD. We then inoculated 200 ul 1.25X SC-Ura + 16 ug/mL Cytosine media with 50 ul resuspended cells, leading to starting ODs between 0.08 and 0.14. Measurements were taken every 15 minutes over 21 hours using a Tecan Infinite M Nano (Tecan Life Sciences) at 30°C. We processed two or three quadrants at a time, allowing us to cover the all-by-all matings in 8 batches of growth curves.

As ODs were not precisely adjusted, there was variation in starting cell density and the lag time. In the case of fast-growing strains (like Fcy1_WT_/Fcy1_LOF_ crosses), this did not impact maximum growth rate measurements, as the curves saturate after 12 to 15 hours.

Measurements for slow-growing strains, which can take longer to attain maximum growth rate, were more affected by this bias(Figure S11a). This explains the lower correlation between cross replicates with intermediate levels of growth. We excluded two samples, MATa Fcy1_N39F_/ MATα Fcy1_R136I_ due to irregularities in the optical density measurements and MATa Fcy1_M93G_/ MATα Fcy1_M93G_ due to cross-contamination with another genotype in the diploid strain stock. The average of cross-replicate measurements was used for downstream when both cross directions were successfully generated (779/1,179 allele pairs). Based on the distribution of growth rates, we defined 0.02 as a conservative threshold for growth (Figure S11b). Growth rates for each allele combination is shown in table S8.

We explored two different models to predict the expected growth rate of an allele pair based on the measured growth rates of heterozygous deletion mutants of both Fcy1 variants. First, we used a dominance (best of both alleles) model where the growth rate of a strain corresponds to the maximum rate measured between the two alleles present in the strain.

Second, we used an additive model, where the expected growth rate is the average of both alleles. Since all crosses with the wild-type allele grew at the same rate independent of the function level of the other allele (Figure S11c), we concluded that the most appropriate model was the one based on dominance. We defined a *trans* epistatic interaction as a growth rate measurement deviation from the dominant model expectation by more than 4 standard deviations of the wild-type cross distribution. This resulted in 164/684 (24.0%) alleles pairs passing the threshold for a *trans* epistatic interaction at an expected false positive rate of less than 0.05%, meaning that the false discovery rate should be well below 1% (0.00064/0.24 = 0.0027%)

To make dose-response measurements, one cross-replicate per chosen mutant was selected at random as a representative in the experiment. Diploid mating glycerol stock plates were thawed on ice, and strains were revived by spotting 5 ul of the stock on YPD+Hyg+Nat and incubating at 30 °C for 48h. Strains were sequentially arranged in the order they appeared in the randomized mating arrays. Each spot was picked twice to inoculate two replicate pre-cultures of each strain in 600 μL SC (MSG, pH buffered)+Hyg+Nat media. The next morning, cells were spun down and adjusted to 0.4 OD in sterile water. We then added 20 μL cells to 60 μL 1.33X SC -Ura + cytosine (either 12, 16, 20 or 24 μg/mL) in a 384-well, resulting in 1X media at 0.1 OD. Optical density was measured every 15 minutes for 48 h without agitation on a Tecan Spark (Tecan Life Sciences) with an active Tecool temperature control module. To limit the effect of water evaporation, the border wells of the plate were filled with sterile water. For each strain, we measured the maximum growth rate over 40 hours. For each compensating allele pair, we tested if its growth rate was significantly lower than the ancestor strain using a one-sided Welch’s t-test. This was repeated for across all cytosine concentrations. If the p-value was above 0.1, we considered growth rates to be equal.

### Haploinsufficiency dose-response assays

Starting from strains s_005 and s_006, we used our usual CRISPR knock-in approach to generate multiple independent *FCY1*_codon opt_ strains in both mating types (s_007 and s_008). We then mated these strains together to generate 4 different homozygous wild-type strains(*FCY1*_codon opt_/*FCY1*_codon opt_). We also crossed two MATa and two MATα with the opposite deletion mutant to generate 4 independent heterozygous mutants (2 x *FCY1*_codon opt_/*fcy1Δ::stuffer3* and 2 x *fcy1Δ::stuffer3*/FCY1_codon opt_). We started 5 mL overnight SC complete (MSG, pH buffered) + Nat + Hyg precultures for each of those strains. The next morning, the cells were diluted to 2 OD in sterile water and then used to inoculate 190 μL SC - Ura + Cytosine 1.05X media in a 96-well plate. The experiment included 12 different cytosine concentrations (0, 4, 6, 7, 8, 9, 10, 12, 16, 20, 24 and 30 μg/mL. Measurements were taken every 15 minutes over 48 hours using a Tecan Infinite M Nano (Tecan Life Sciences) at 30°C. To determine the IC_50_, we fit a Hill equation to the the relative maximum growth rates over the first 24h, using the average growth rate of the homozygous strains at 30 μg/mL as the reference for maximal response.

### Inducible promoter haploid strains

First, to be able to combine two single mutants in an haploid cell, we deleted *FCY1* with *HphNT1* in the landing pad strain (s_096, BY4741 *leu2::GEM Δfcy1::HphNT1 Δgal1::stuffer3*). Then using two consecutives CRISPR-Cas9 transformation rounds targeting either *HphNT1* or *stuffer3*, (Ryan et al 2016) we introduced three *FCY1* alleles at each position (wild-type, E64V and R73G) to get all possible combinations. We also kept the intermediate strains where there was only one allele present at either loci, for a total of 15 strains. We selected strains s_109 (*leu2::GEM FCY1_R73G_ Δgal1::FCY1*_E64V_), s_098 (*Δfcy1::HphNT1 Δgal1::FCY1*_E64V_), s_110 (*leu2::GEM FCY1*_E64V_ *Δgal1::FCY1*_R73G_) and s_099 (*Δfcy1::HphNT1 Δgal1::FCY1*_R73G_) to compare growth based on the induced allele.

We started 3 independent cultures of each strain in SC -leu (MSG) and let it grow overnight. The next morning each culture was diluted to 0.15 OD in fresh SC -leu (MSG) with 4 different concentrations of beta-estradiol (0, 4.5, 6 and 16 nM in ethanol). Once cultures reached ∼0.7 OD/ml, cells were spun down and washed in water to a final OD of 1. Then 25 µl of cells were transferred to 225µl of two different media : SC -leu (MSG) or SC -leu -ura (MSG) +16 µg/ml cytosine making sure to keep the same concentration of inducer as in the intermediate culture. OD was read each 15 min at 30°C for 2 days without shaking.

### Generation of *FCY1* double mutant alleles in cis and growth assays

To systematically explore a small space of double mutant alleles, we used the same fusion PCR approach as described above, but using *FCY1* variant alleles as template instead of the wild-type sequence. Because we used the same oligonucleotides as previously, some strains were impossible to generate as the template mutation fell into the window covered by the mutagenesis oligo (e.g.: *FCY1_H62C,E64V_*). Single mutant control alleles for this experiment were constructed by using the same oligonucleotides used to generate the template strain again. We inserted these alleles in s_005 (BY4741 *fcy1Δ::stuffer3 ΔHO::NatNT2)* via a CRISPR transformation as described above. We then validated strain construction via Sanger sequencing, resulting in a set of strains covering 48/64 pairwise mutant permutations and 33/36 mutant combinations.

We then tested the growth of these strains in 5-FC and cytosine media. From the rearrayed strains, two pre-cultures were inoculated per strain in 500 μl SC complete (MSG, pH buffered) + Nat for overnight growth. The next morning, cells were spun down and adjusted to 0.4 OD in sterile water. We then added 20 μL of cells to either 60 μL of 1.33X SC + 5-FC (100 μg/mL) or 60 μL 1.33X SC -Ura + cytosine (16 μg/mL) in a 384-well plate, resulting in 1X media at 0.1 OD. Optical density was measured every 15 minutes for 48 h without agitation on a Tecan Spark (Tecan Life Sciences) with an active Tecool temperature control module. To limit the effect of water evaporation, the border wells of the plate were filled with sterile water. For each strain, we measured the maximum growth rate over 40 hours.

To determine whether double mutant FCY1 alleles can maintain the potential for *trans* epistasis, we mated the MATa double mutant array to two other alleles, MATα *FCY1_E64V_* and MATα *FCY1_R73G_*. Briefly, we inoculated 500 μL YPD with 20 ul MATa and 20 μL MATα overnight precultures and incubated at 30C for ∼8 hours before spotting on YPD+Nat+Hyg. We then inoculated two pre-cultures per spot in 500 μL SC complete (MSG, pH buffered) + Nat+Hyg for overnight growth. We then diluted these strains to inoculate growth curves in cytosine media SC -Ura + cytosine (16 μg/mL) in a 384-well plate as described above.

### *FCY1* protein fragment complementation assay

The interaction between Fcy1 dimer subunits can readily be detected by *in vivo* protein complementation assay, DHFR-PCA (*16*, *32*). To generate N-terminal tagged variant alleles, we used a fusion PCR approach similar to our previous work (*11*). We first amplified both DHFR fragments from genomic DNA of strain s_167 (using oligo_238) and s_168 for (using oligo_239) for DHFR F[1,2] and DHFR F[3] respectively. The amplicons were flanked in 5- prime with homology the the *FCY1* promoter and with the DHFR linker in 3-prime (GGGGS repeats). In parallel, we amplified wild-type or mutant FCY1 alleles from their respective genomic DNA using o_237 and o_002, which resulted in amplicons with homology to the linker in 5-prime and homology to the FCY1 terminator in 3-prime. We then fused these amplicons in a second PCR reaction, and used these tagged alleles as donor in a CRISPR transformation using our standard protocol. We transformed MATa strains with the DHFR[1,2] tagged alleles and MATα strains with the DHFR[3] tagged alleles.

Knock-in strains were validated by Sanger sequencing, which revealed a duplication in the linker sequence of DHFR[3]-Fcy1_E64V_ that increased the number of GGGGS repeats from 2 two three. As increased linker length is not detrimental to protein-protein interaction detection by PCA (*33*), we decided to use the strain nonetheless. MATa DHFR F[1,2] strains were crossed with MATα DHFR F[3] strains using the same protocol previously described.

We then performed growth curves in PCA media to detect protein-protein interactions. Diploids were grown overnight in 500 μl SC (MSG) pH buffered+Nat+Hyg. The next morning, cells were diluted to 0.4 OD and used to inoculate 60 ul PCA media with or without MTX (see recipes) in a 384-well plate as described above. Measurements were taken every 15 minutes for 40 hours.

### Fcy1 expression and purification

All heterologous expression was performed in *E. coli* strain BL21 (DE3) (*34*) and followed the same approach with varying culture volumes depending on the scale of the downstream purification step. The coding sequence of Fcy1 (wild-ype or matching mutants of interest) was synthetized in pET28b(+), adding a His-tag (6X) and thrombin cleavage site inserted inframe upstream of the coding sequence (*35*). These plasmids were transformed in freshly prepared CaCl_2_ chemically competent BL21 (DE3) cells using an inhouse protocol. After transformation, 5-10 colonies per transformation plates were picked and used to inoculate a 2X Yeast Tryptone Broth (2YT) + kanamycin 50 μg/mL overnight pre-culture. The next day, this preculture was used to generate glycerol stocks and then was diluted 1:100 in Terrific Broth (TB) media + kanamycin 50 μg/mL. Glycerol stocks were freshly thawed and streaked before inoculating pre-cultures when not working with fresh transformants.

After dilution in TB in flasks, cells were incubated at 37 C with shaking (250 RPM). When cultures reached 0.6-1.0 OD (which usually took 2-3 hours), we added IPTG for expression induction to a concentration of 0.5 mM and zinc acetate (a Fcy1 cofactor) to 1 mM. The cells were then switched to 16°C and incubated overnight (∼16h). The next day, cells were spun down in batches at 3000 RPM, 4°C. The resulting cell pellets were then frozen at -20 until further use.

To co-express two different FCY1 variants in the same *E. coli* cells, we switched the markers of the pET28b(+) from kanamycin to ampicillin using Gibson cloning. First, we amplified the ampicillin marker from pAG25 using CLOP288-F4 and H4. In parallel, we amplified the pET28b(+) backbone with CLOP288-F2 and G2. Template DNA was removed with DpnI treatment for 1h at 37°C and then we purified both PCR products using magnetic beads.

After purification, the amplified backbone and resistance marker were mixed at a 1:3 ratio in in-house Gibson assembly mix and incubated 1h at 50°C. Following the incubation the Gibson assembly was transformed into bacteria, plated on 2YT+AMP petri dish and grown overnight at 37°C. Four colonies were selected for each cloning reaction and plasmids were extracted using the PRESTO miniprep kit (Geneaid, Taiwan). To confirm integration of the ampicillin marker plasmid were digested with ScaI and XbaI (NEB R3122L and R0145L) and verified using gel electrophoresis. Clones showing the predicted restriction profile and able to growth in 2YT+AMP were considered positives. Plasmids were either co-transformed at the same time or in a second round of transformation where competent cells were generated from one of the plasmid bearing strains. We used the same induction protocol during co-expression, but adding ampicillin 100 μg/mL in addition to Kanamycin in the pre-culture and induction media.

### Fcy1 homomer affinity purification

We used two approaches for affinity based purification of Fcy1 after heterologous expression in *E. coli* depending on the scale required. Small-scale purification was used to purify proteins for enzymatic activity measurement, while large-scale purification was used to produce large quantities of Fcy1 for crystallography and stability measurements.

For small-scale purification, we used cell pellets equivalent to 25 mL of post-induction cell culture. The pellets were resuspended in 1.25 mL of NEB quick lysis buffer (NEB, Ipswich, USA) supplemented with 0.5 mg/mL lysozyme, 0.17 mg/mL phenylmethylsulfonyl fluoride (PMSF) in isopropanol and 0.01 mg/mL DNAse I from 1000x stocks. Cells were lysed for 10 minutes at room temperature on a rotary shaker. The resulting cell lysate was centrifuged at 21,300 g for 15 minutes at 4C. We then used the Dynabeads His-Tag Isolation and Pull-down kit (ThermoFisher 10103D) to isolate his-tagged Fcy1 following manufacturer’s instructions. Briefly, we mixed 700 μL cell lysate and 50 μL magnetic beads and incubated for 10 minutes on a rotary mixer to allow the tagged proteins to bind. The beads were then washed 4 times with cold washing buffer (50 mM sodium-phosphate pH 8.0, 300 mM NaCl, 0.01% Tween-20), leaving the tubes two minutes on a magnetic rack to let the pellets form. The purified proteins were then eluted by resuspending the beads in elution buffer (300mM imidazole, 50 mM sodium-phosphate pH 8.0, 300 mM NaCl, 0.01% Tween-20), incubating for 5 minutes on a rotary roller, and collecting the supernatant. Right after elution, we performed a buffer-exchange to PBS (9.55 mM Na_2_H_2_PO_4_,137 mM NaCl, 2.65 mM KCl, 1.47 mM KHPO_4_) + 0.01% Tween-20 using Zeba 0.5 mL Spin Column 7K (ThermoFisher 89882). The resulting purified Fcy1 retains the N-terminal His-tag and thrombin cleavage site, and were used fresh for enzymatic assays. Example Coomassie gels for Fcy1_WT_, Fcy1_E64V_ and Fcy1_R73G_ small scale affinity purification are shown in Figure S17.

For large-scale purification, we used cell pellets equivalent to larger culture volumes (400 mL, 800 mL or 1 L). The cells were resuspended in inhouse lysis buffer (50 mM Tris-HCl pH 7.5, 400 mM NaCl, 2% glycerol, 5mM Imidazole, 0.5 mg/mL lysozyme, 0.17 mg/mL PMSF, 0.01mg/mL DNAse I) and lysed by sonication (10 minutes, 35% amplitude in 10 s on, 20 s off cycles). The cell lysate was then spun at 13 000 rpm for 50 minutes at 4°C before being loaded on a Cobalt ion resin and incubating for 45 minutes at 7°C. We then washed the column 3 successive times using the following buffers with 5 minutes incubation: 50 mL lysis buffer, 25 mL washing buffer 2 (50 mM Tris-HCl pH 8.0, 1 M NaCl, 5% glycerol, 10mM Imidazole) and 25 mL washing buffer 3 (50 mM Tris-HCl pH 8.0, 400 mM NaCl, 5% glycerol, 15 mM Imidazole). The purified proteins were then eluted in two fractions using 3 mL (50 mM Tris-HCl pH 7.5, 1 M NaCl, 5% glycerol, 200 mM Imidazole) with 10 minutes incubation. After elution, Fcy1 was exchanged to a protein buffer containing 20 mM Tris-HCl pH 7.5, 150 mM NaCl, and 2% glycerol. The His-tag was cleaved by adding 7 U of thrombin per milligram of protein and the resulting cleaved protein (passing through the Ni-NTA beads) was concentrated to 15 mg/ml. Purification of the Fcy1_M100W_ mutant was achieved using the same method as described above.

### Fcy1_E64V_-Fcy1_M100W_ heteromer affinity purification

The coding sequence of Fcy1_E64V_ was synthesized in pET28a(+), adding a Strep-tagII© (IBA Lifesciences GmbH) and thrombin cleavage site inserted in frame upstream of the coding sequence. This plasmid was co-transformed with the pET28b(+) plasmid of Fcy1_M100W_ with a N-terminal 6-His-tag and thrombin cleavage site which was modified to have an ampicillin marker instead of kanamycin to yield a heterodimer with different tags. Expression was done as stated before. Theoretically speaking, three products can be obtained: the StrepII-tagged homodimer of Fcy1_E64V_, the His-tagged homodimer Fcy1_M100W_ and the heterodimer of StrepII-tagged Fcy1_E64V_ and His-tagged Fcy1_M100W_. We aimed to use tandem affinity chromatography to purify the heterodimer as it would be the only species with both affinity tags.

We first purified the heterodimer and the StrepII-Fcy1_E64V_ species by StrepII-tag affinity. We used a cell pellet from 1.6 L of culture resuspended in lysis buffer (50 mM Tris-HCl pH 8.0, 400 mM NaCl, 2% glycerol, 5mM Imidazole, 0.5 mg/mL Lysozyme, 0.17 mg/mL PMSF, 0.01mg/mL DNAse I) and then did sonication (10 minutes, 35% amplitude in 10 s on, 20 s off cycles). The cell lysate was centrifuged at 13 000 rpm for 30 minutes at 4°C. Strep-tagged proteins were captured on Strep-Tactin XT Superflow High Capacity (IBA LifeSciences) affinity resin. Following a washing step (150 mM NaCl, 50 mM Tris-HCl pH 8.0), the proteins were eluted in 1x BXT buffer (150 mM NaCl, 50 mM Tris-HCl pH 8.0, 1mM EDTA; IBA LifeSciences). We then exchanged the buffer to the protein buffer containing 20 mM Tris-HCl pH 8.0, 150 mM NaCl, and 2% glycerol and captured the heterodimer species on cobalt resin. The sample was incubated 20 minutes at 4°C and was washed with a gradient of imidazole, then eluted in elution buffer (50 mM Tris-HCl pH 7.5, 1 M NaCl, 5% glycerol, 200 mM imidazole). We used buffer exchange to obtain the purified heterodimer buffer containing 20 mM Tris-HCl pH 8.0, 150 mM NaCl, and 2% glycerol. The heterodimer was concentrated to 12 mg/ml before crystallization trials.

### Crystallization of wild-type Fcy1, Fcy1_M100W_ mutant and Fcy1_E64V_-Fcy1_M100W_ heteromer

Multiple crystallization screening kits from NeXtal Biotechnologies (Holland, OH, USA) were used to get the initial hits. Two crystal forms of wild-type Fcy1 were obtained using the microbatch-under-oil method in distinct reservoir solutions: (a) 0.1 M sodium cacodylate pH 6.0, 15% PEG-4000; (b) 0.2 M lithium sulfate, 0.1 M MES pH 6.0 and 20% PEG-4000.

Crystals of the Fcy1_M100W_ mutant were harvested from the reservoir solution containing 10% 2-Propanol, 0.1 M HEPES pH 7, and 10% PEG 4000. The Fcy1_E64V_-Fcy1_M100W_ heteromer was crystallized from the reservoir solution containing 0.2 M Ammonium chloride, 0.1 M MES pH 6, and 20% (w/v) PEG 6000.

### Crystallographic data collection and structure determination

Both data sets of wild-type Fcy1 were collected at the LRL-CAT beamline at the Advanced Photon Source, Argonne National Laboratory. The anomalous data sets for two crystals obtained in the presence of sodium cacodylate were collected at the wavelengths of 1.04339 Å and 1.27898 Å, respectively while the anomalous data set for the crystal obtained in the presence of lithium sulfate was collected at the wavelength of 1.27899 Å. Datasets for the Fcy1_M100W_ mutant and the Fcy1_E64V_-Fcy1_M100W_ heterodimer were collected at the wavelength of 0.95374 Å at the CMCF-ID beamline at the Canadian Light Source.

Data processing and scaling were performed with XDS (*36*). Both Fcy1_WT_ crystals were in the P21 space group. Those obtained in the presence of cacodylate have unit cell parameters of a=39.11, b=55.07, c=66.87 Å, β=90.5° while the crystals grown in the presence of lithium sulfate have cell dimensions of a=77.86, b=40.83, c=97.73 Å, β=104.8°. The Fcy1_M100W_ mutant was crystallized in the P1 space group with a =79.9 b =81.4, c =168.2 Å, α =89.8°, β =90.2°, γ= 90.8° while the crystal of the Fcy1_E64V_-Fcy1_M100W_ heterodimer was in the P212121 space group with a = 46.9, b =77.7, c =148.8 Å. The structures were determined by MOLREP (*37*) in CCP4 (*38*) using the template previously published (PDB 1OX7). To get the final structures, multiple cycles of refinement using Refmac5 (*39*) followed by manual model correction with Coot (*40*) were carried out. The stereochemistry of all the models reported here has been analyzed with PROCHECK (*41*). Data collection and refinement statistics are shown in table S1.

### Active site cofactor reannotation

Previous structural studies on Fcy1 (*12*, *42*, *43*) have indicated the presence of two Zn ions, one catalytic and the other noncatalytic, in the active site of each Fcy1 subunit in the absence of substrate or inhibitor. The tetrahedral coordination sphere of the catalytic Zn ion includes the ND1 atom of His62, the thiol groups of both Cys91 and Cys94, and one water molecule while the noncatalytic Zn is tetrahedrally coordinated by four water molecules including the one bound to the catalytic Zn. Of note, no protein atom is in the coordination sphere of noncatalytic Zn and two water molecules are even less than 1.8 Å distant from this Zn ion in the best resolution (1.43 Å, PDB 1OX7) model of such Fcy1 structures.

Nonetheless, a survey of the Zn binding environment in a large number of protein structures indicated the distance between Zn and the surrounding water molecules is between 2.1 and 2.3 Å (*44*). This discrepancy might imply that this noncatalytic Zn is in fact not a Zn ion but rather another small molecule captured in the active site. Indeed, cacodylate in the reservoir solution could be a good candidate as it fits well into the electron density at this location given its tetrahedral geometry for the oxygen and carbon atoms covalently linked to the arsenic atom with a bond length of 1.7 ∼ 1.8 Å (As-O) or 1.9 ∼ 2.0 Å (As-C).

To confirm our hypothesis, we crystallized wild-type Fcy1 and have obtained two crystal forms in different solutions. Similar to the crystallization conditions previously used, the first crystal form was obtained in the presence of cacodylate. Each asymmetric unit contains a biological dimer. Except a few residues at the N-terminal, the structure is almost identical to all the available Fcy1 structures previously determined (obtained in the presence of cacodylate or inhibitor), as exemplified by a root mean square deviation of 0.27 Å for the superposed 302 Cα atoms between our Fcy1 dimer and the structure 1OX7 (*12*). The anomalous difference map calculated from the 1.39 Å resolution data collected at As edge (a wavelength of 1.04339 Å) reveals strong peaks for both As (38.7σ and 36.8σ) and catalytic Zn (26.8σ and 26.6σ) atoms in the active sites of both subunits, strongly supporting the presence of cacodylate in the active site. Moreover, anomalous difference map calculated from the 2.03 Å resolution data collected for a second isomorphous crystal grown in the presence of cacodylate at the wavelength of 1. 27898 Å (Zn edge) shows only one strong peak for each subunit (16.2σ and 13.1σ for subunits A and B, respectively) corresponding to the location of the catalytic Zn, unambiguously confirming the presence of only one Zn atom in the active site.

### Fcy1_WT_ K_D_ determination

To determine the Fcy1_WT_-Fcy1_WT_ binding K_D_, we measured emission spectral shift of covalently fluorescent labeled Fcy1_WT_ using a Monolith X platform (NanoTemper, Germany). We used the Monolith Protein Labeling Kit RED-NHS 2nd Generation kit (NanoTemper MO- L011) to label ∼1 lysine per Fcy1 monomer with a fluorophore following the manufacturer’s instruction. Briefly, 10 ug Dye was reconstituted in 25 ul DMSO (final concentration 600 uM), diluted 1:1 (7ul : 7ul) in labeling buffer. We then added 10 ul 300 uM dye to 90 ul 10 uM Fcy1 in PBS-T, mixed thoroughly, and incubated the mixture at room temperature in the dark for 30 minutes. After labeling, the remaining dye was removed via buffer exchange using the provided B-column equilibrated with PBS-T.

To perform the spectral shift assay, we mixed different ratios of Fcy1_Labeled_ and Fcy1_Unlabeled_ following the AssayControl software (NanoTemper, Germany) suggested protocol. This resulted in a 16 point two-fold dilution series of Fcy1_Unlabeled_ spanning from 5 uM to 150 pM. For each dilution, 10 ul was sampled in a Monolith.NT115 capillary (NanoTemper MO-K022) and measurements were performed at 25°C using standard parameters. We excluded one dilution (4.88 nM) across all measurements as an outlier. Our K_D_ measurement assay followed best practices in measuring binding affinities (*45*). First, to ensure K_D_ was measured at equilibrium, we performed the spectral shift assay at multiple timepoints over 2.5 hours with the same capillaries, ending the experiment when the K_D_ was stable and the signal to noise ratio started dropping. Second, the concentration of Fcy1_Labeled_ (20 nM) is only ∼4 times the K_D_, which is safely within the concentration range associated with a binding regime. To estimate the cellular concentration of Fcy1, divided its consensus abundance of 8488 molecules/cell (*46*) by the median volume of a haploid yeast cell, 42 μm^3^ (BNID 100427, (*47*)). To obtain the fraction of Fcy1 molecules bounds as dimers *in vivo*, we then used the protein homodimerization simulator from (*48*) with the monomer molecular mass, the measured K_D_ and the estimated cellular concentration as input.

### Fcy1 variants stability and activity testing

We measured protein stability using nano-differential scanning fluorimetry on a Prometheus NT.48 (NanoTemper). After purification and his-tag cleavage, 10 ul protein sample was loaded in a Prometheus NT.48 Series nanoDSF Grade Standard Capillaries (NanoTemper PR-C002). Measurements were taken along a 25°C to 95°C temperature gradient, increasing at a rate of 1C/min. The T_m_ was measured by finding the temperature at which the first derivative of the 350/330 nm fluorescence ratio is maximised using the Thermo Control software v2.3.1. Each protein variant was tested in triplicate in the same experiment, and the median T_m_ was reported (table S9).

We measured Fcy1 activity by following the conversion of 5-FC into 5-FU by measuring the shift in absorbance at 237 nm in a Corning UV transparent 96-well plate (ref# 3635). For each assay, we mixed 100 μL 0.0125mg/ml purified Fcy1 with 100 μL 0.225 mg/mL 5-FC in triplicate and then measured absorbance every 90 seconds over 3 hours. All measurements were performed in a Tecan Spark (Tecan Life Sciences) with an active Tecool temperature control module set at 25°C. Measurements were first scaled to represent the fraction of 5-FC converted relative to the last wild-type timepoints, and then to the activity of the wild-type to normalize for batch effects.

## Data and code availability

Raw sequencing file have been deposited on the NCBI SRA (PRJNA1061126). Protein structures have been deposited with the RCSB PDB: Fcy1-closed: 8VLJ, Fcy1-open: 8VLK, Fcy1_M100W_: 8VLL, Fcy1_E64V_-Fcy1_M100W_: 8VLM. Source data for data analysis and figure generation are available as supplementary material. The scripts used are available at https://github.com/Landrylab/Despres_et_al_2024. The analysis scripts made use of the following python packages: numpy (*49*), scipy (*50*), matplotlib (*51*), pandas (*52*), seaborn (*53*) and Biopython (*54*). Analysis scripts also used the following software: fastqc (*55*), Trimmomatic (*56*), PANDAseq (*30*) and vsearch (*31*).

## Acknowledgements

The authors thank D. Evans-Yamamoto, M. Hénault, F. Mattenberger, S. Dibyachintan and S. Gobeil and Steve Michnick for their insightful comments and discussion around this manuscript. This work was supported by the Canadian Institutes of Health Research Foundation grant number 387697 to C.R.L. and a Vanier graduate scholarship to P.C.D, the National Science and Engineering Research Council through the EvoFunPath CREATE grant (555337-2021) to P.C.D., J.G. and C.R.L. and a Discovery Grant (436202) to R.S., FRQNT through team grant (number 2022-PR-298169) to R.S. and C.R.L., and FRQS through a Doctoral training award to P.C.D. C.R.L. holds the Canada Research Chair in Cellular Systems and Synthetic Biology.

Part of the research described in this paper was performed using beamline CMCF-ID at the Canadian Light Source, a national research facility of the University of Saskatchewan, which is supported by the Canada Foundation for Innovation (CFI), the Natural Sciences and Engineering Research Council (NSERC), the National Research Council (NRC), the Canadian Institutes of Health Research (CIHR), the Government of Saskatchewan, and the University of Saskatchewan. This research also used resources of the Advanced Photon Source; a U.S. Department of Energy (DOE) Office of Science User Facility operated for the DOE Office of Science by Argonne National Laboratory under Contract No. DE-AC02- 06CH11357. Use of the Lilly Research Laboratories Collaborative Access Team (LRL-CAT) beamline at Sector 31 of the Advanced Photon Source was provided by Eli Lilly and Company, which operates the facility. Molecular graphics and analyses were performed with UCSF ChimeraX, developed by the Resource for Biocomputing, Visualization, and Informatics at the University of California, San Francisco, with support from National Institutes of Health R01-GM129325 and the Office of Cyber Infrastructure and Computational Biology, National Institute of Allergy and Infectious Diseases.

## Contributions

P.C.D., A.K.D., R.S. and C.R.L. designed research. P.C.D., A.K.D., J.G. and M.-E. P. performed experiments and R.S. performed the crystallographic analysis. P.C.D. analyzed the data and wrote the manuscript with CRL and input from all authors.

## Supplementary Figures 1-17

**Figure S1:**
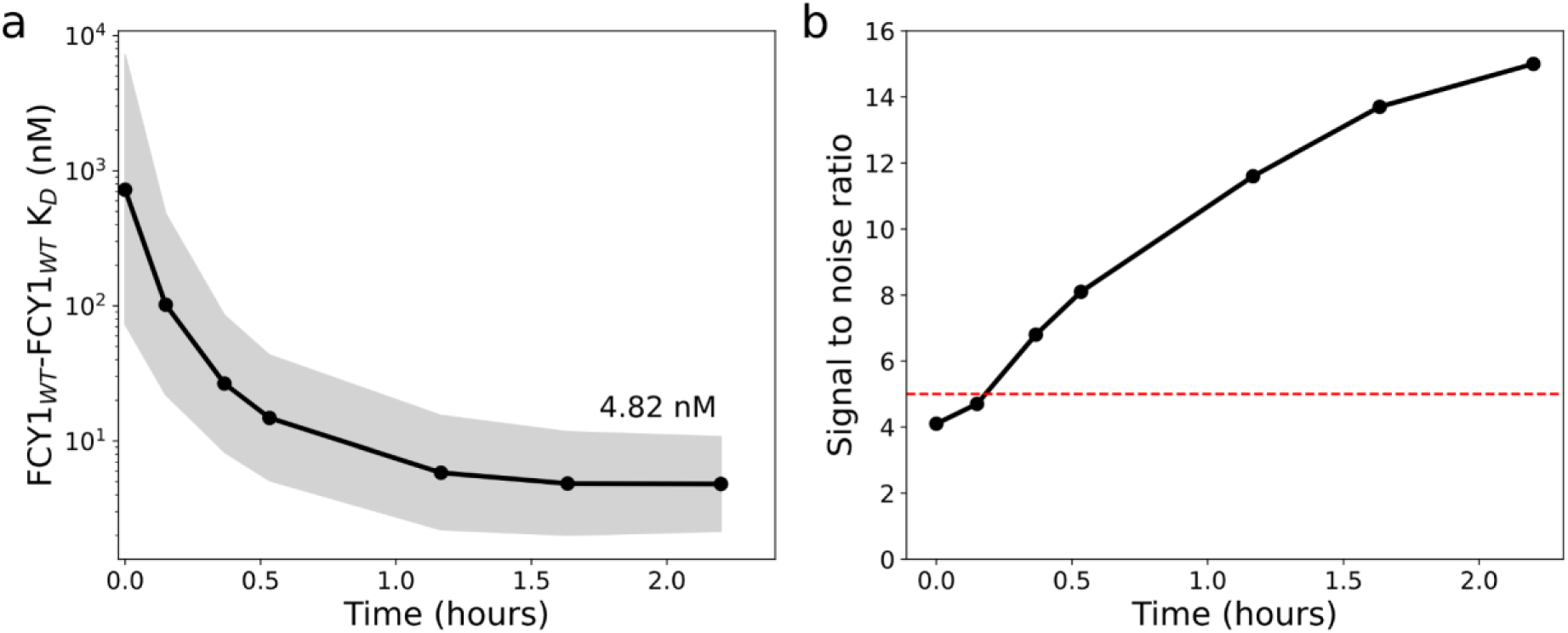
Fcy1 K_D_ measurements show high affinity between dimer subunits. a) Measured K_D_ for the Fcy1-Fcy1 homodimer measured at different time points after mixing samples. The grey area represents the upper and lower bounds of the 95% confidence interval around the K_D_. b) Signal to noise ratio of the binding assay at the different time points. As the sample gets closer to equilibrium, the signal to noise ratio increases.

**Figure S2.**
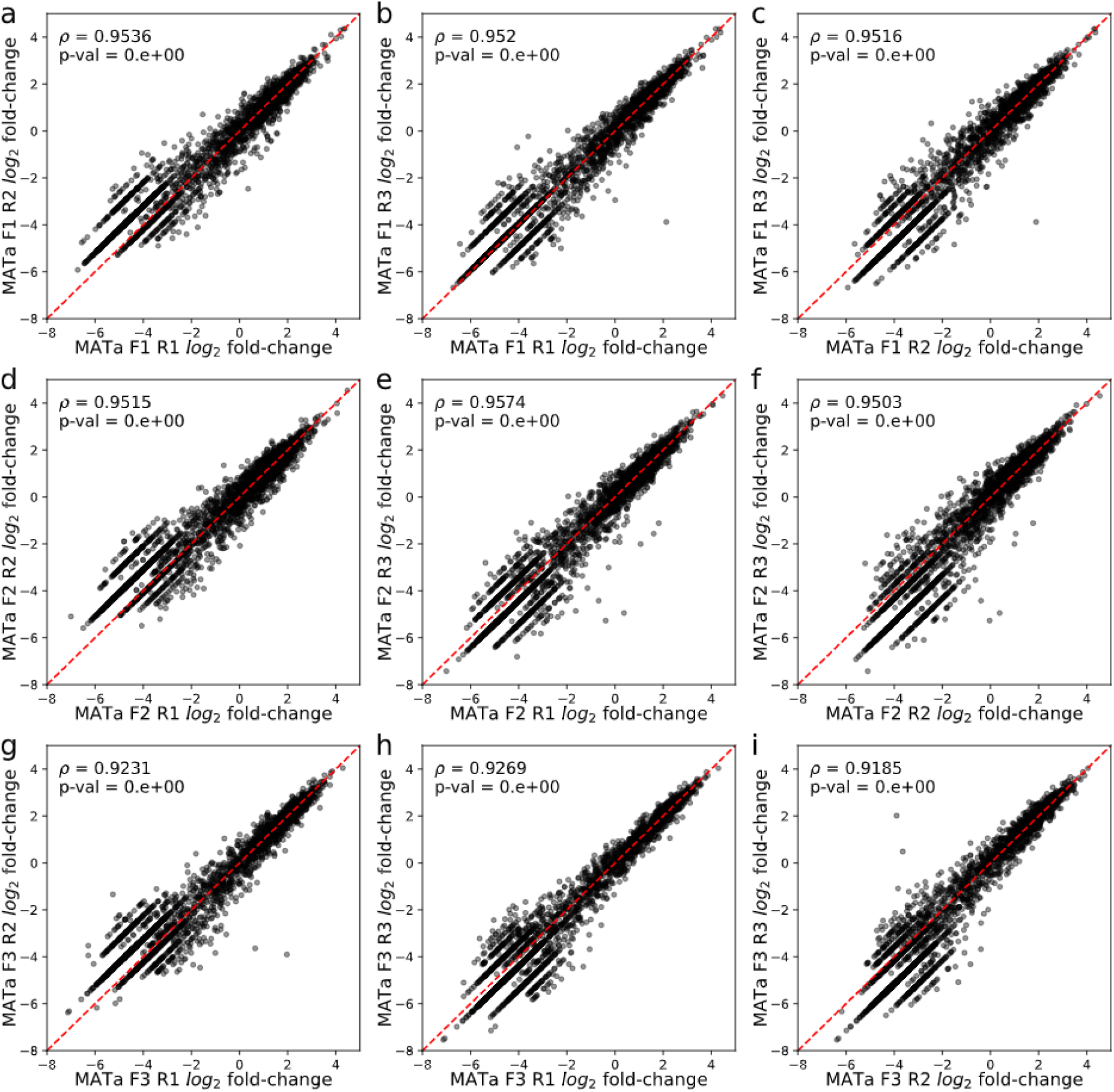
Codon-level *FCY1* allele log2 fold-changes are well correlated between replicates for MATa pools. **a)**, **b)** and **c)** correlation between replicates of fragment 1 (codons 2 to 67, n=4,042). **d)**, **e)**, **f)** correlation between replicates of fragment 2 (codons 49- 110, n= 3,809). **g)**, **h)**, **i)** correlation between replicates of fragment 3 (codons 93-158, n=3,908).

**Figure S3.**
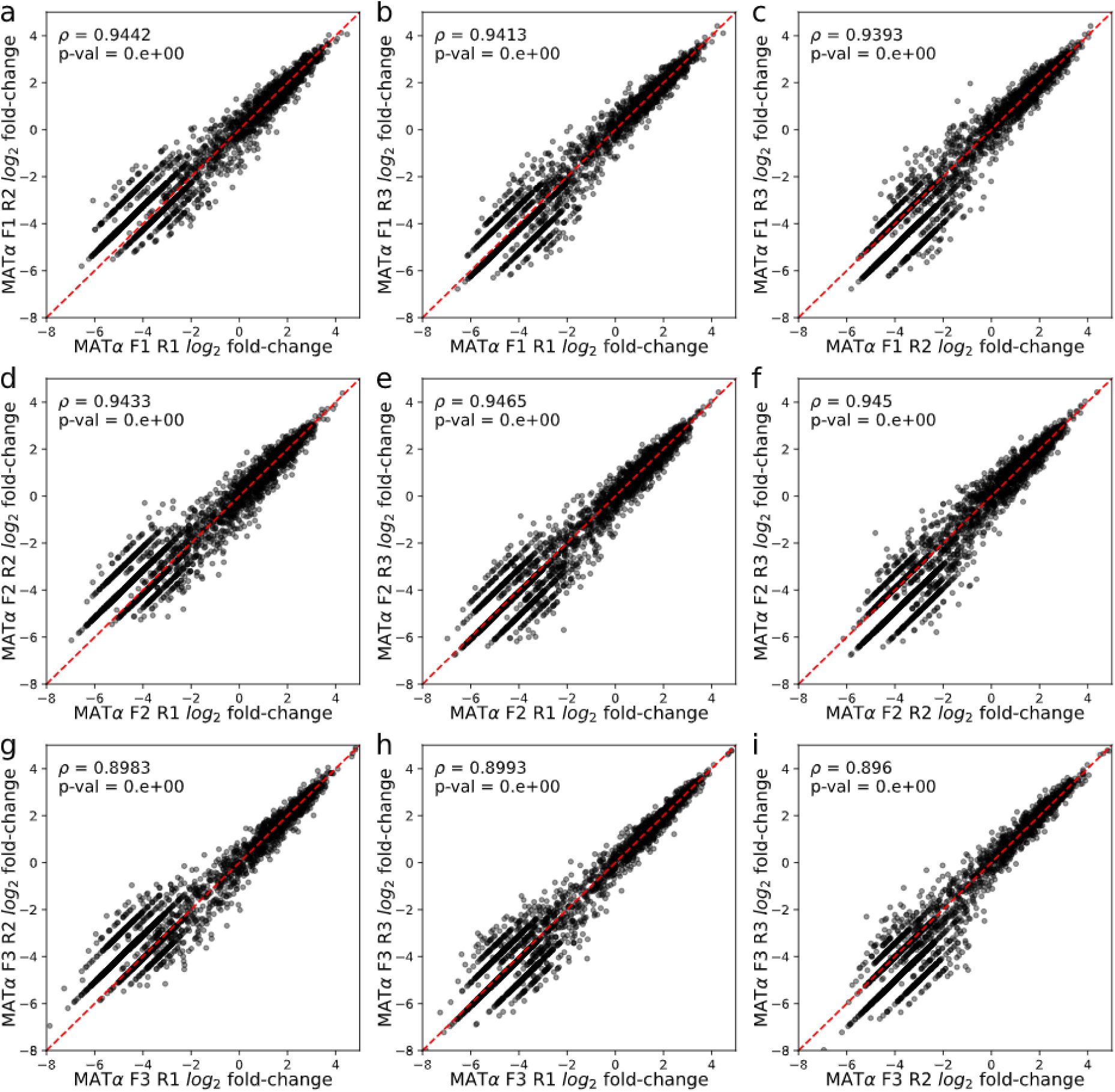
Codon-level *FCY1* allele log2 fold-changes are well correlated between replicates for MATα pools. **a)**, **b)** and **c)** correlation between replicates of fragment 1 (codons 2 to 67, n=4,000). **d)**, **e)**, **f)** correlation between replicates of fragment 2 (codons 49- 110, n= 3,825). **g)**, **h)**, **i)** correlation between replicates of fragment 3 (codons 93-158, n=3,919).

**Figure S4.**
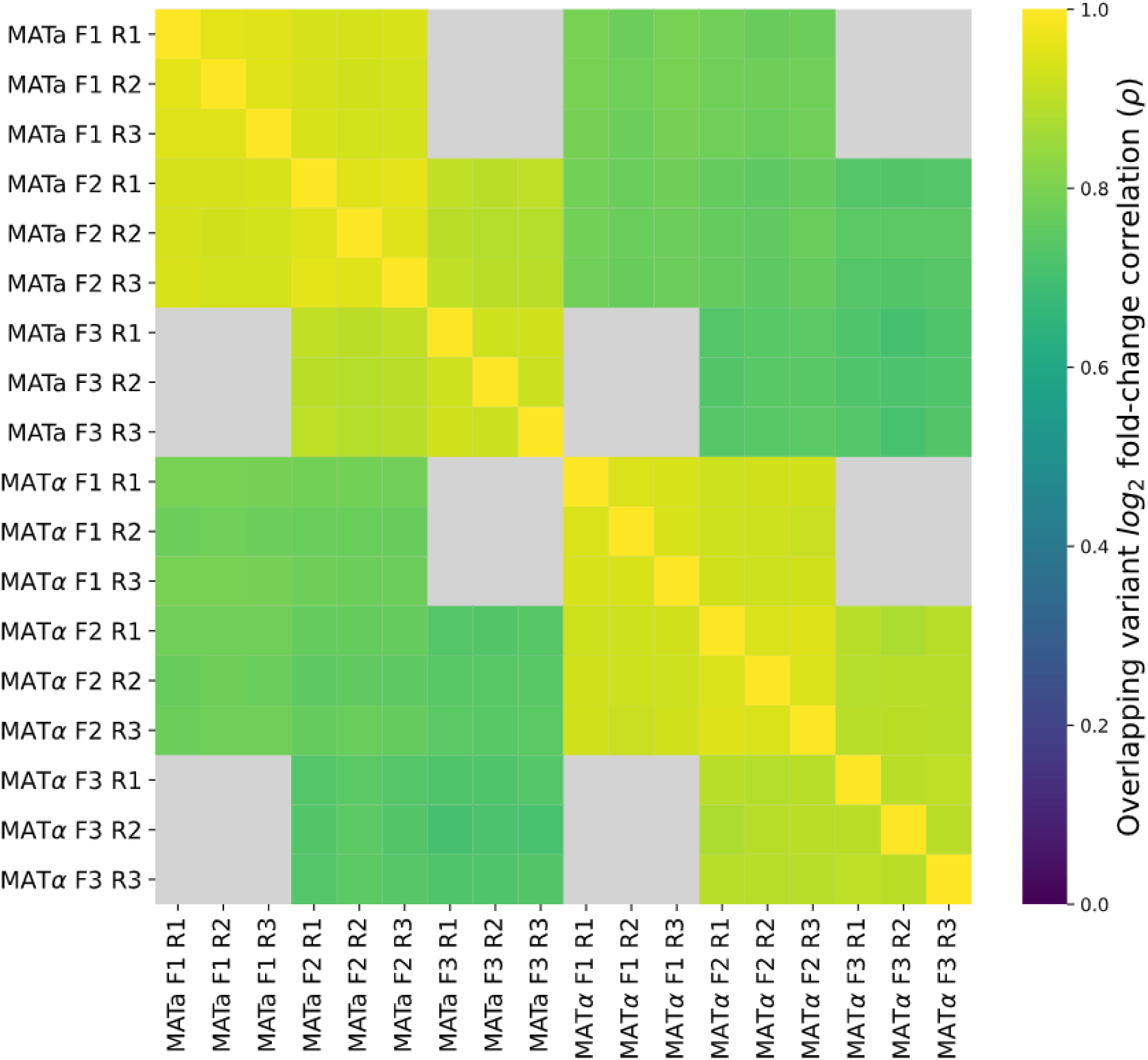
Codon-level *FCY1* allele log2 fold-changes are well correlated across all variant pools after negative selection. Variant pools clustered by Spearman’s rank correlation of log_2_ fold-changes between T0 and the fourth negative selection passage in 5- FC. Grey squares represent Fragment 1-Fragment 3 pairs of sample pools that did not share any variants.

**Figure S5.**
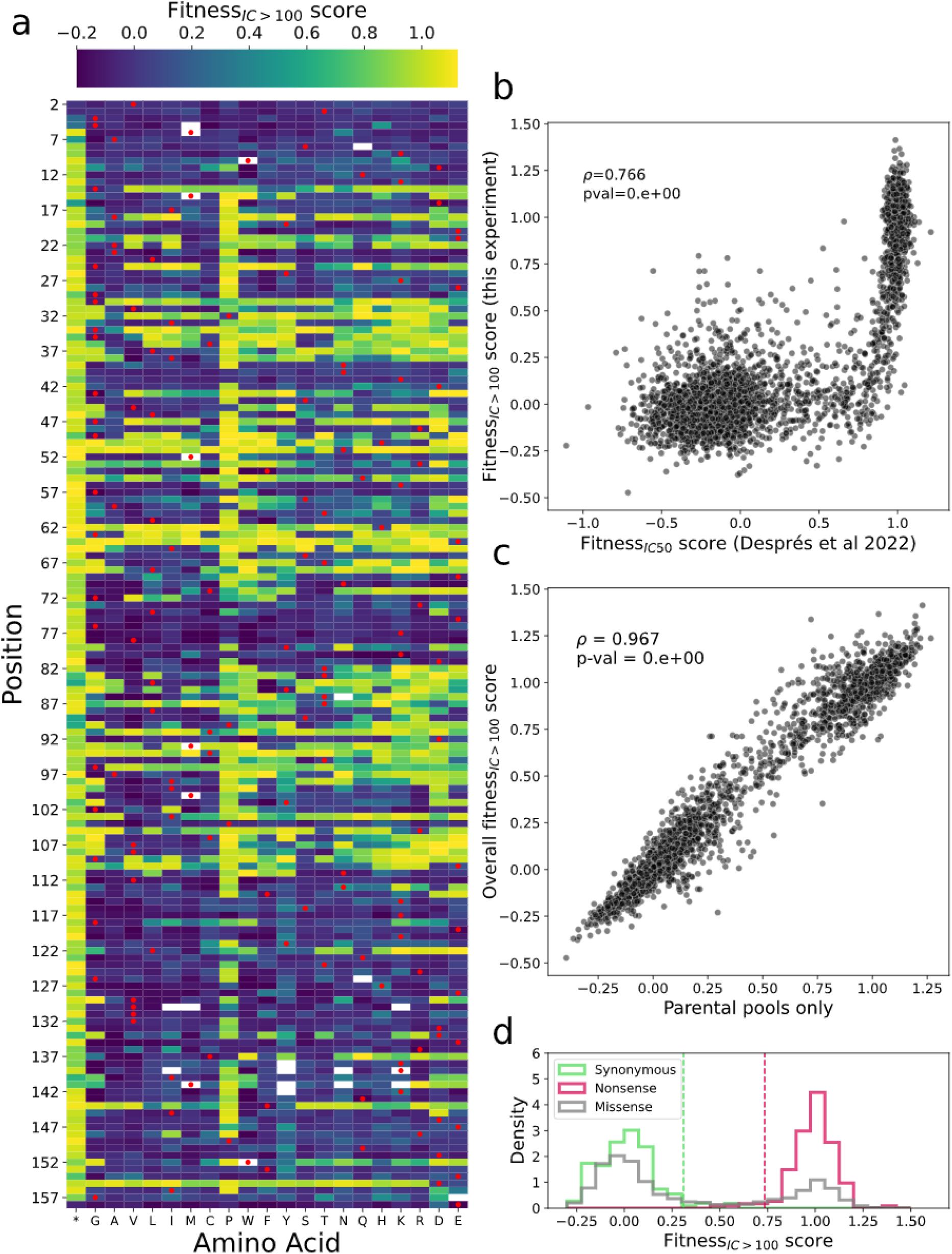
High 5-FC concentrations select for Fcy1 variants with strong LOF effects. **a)** Median fitness score across all pools for single amino acid mutants. Colorbar minimum and maximum correspond to the 2nd and 98th percentile of values. Red dots represent the wild-type amino acid for each position. **b)** Correlation between fitness scores of amino acid level variants (n=3,272, including synonymous and nonsense mutants) observed in this study (IC_>100_, 100 μg/mL 5-FC) and those from our previous assay at a lower 5-FC concentration IC_50_, 1.56 μg/mL (*11*). **c)** Fitness scores for single amino acid variants as measured in the parental pools used to make the LOF x LOF crosses compared with scores obtained across all pools (n=3,272). **d)** Distribution of fitness scores for synonymous, nonsense and missense Fcy1 variants in parental pools. The green dotted line represents the 97th percentile of synonymous mutant scores, which was used as the threshold to classify variants as LOFs. The red dotted line represents the 3rd percentile of nonsense mutant scores.

**Figure S6.**
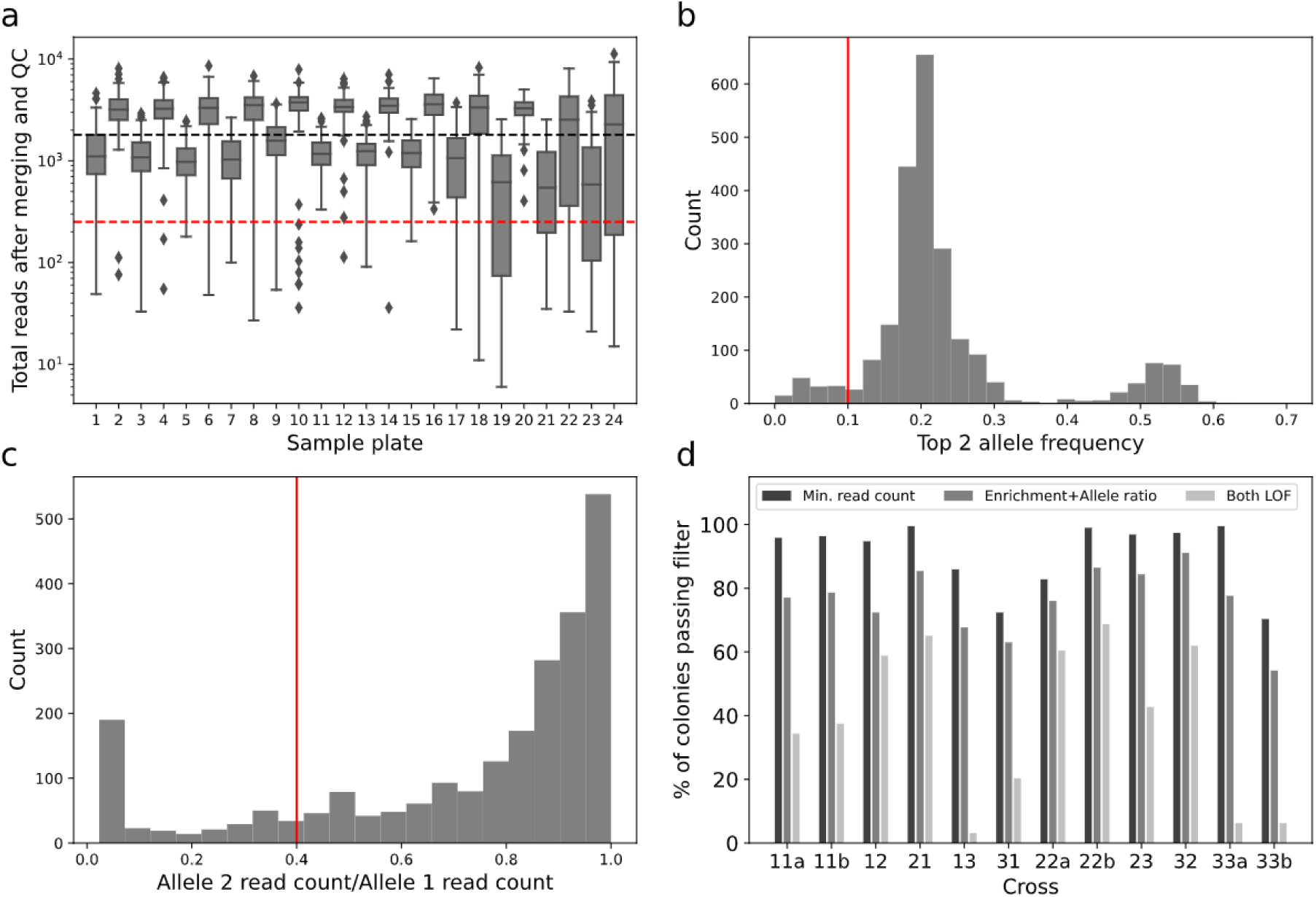
Filtering steps to identify auxotrophy complementing LOF variant pairs. **a)** Read count distribution for each sequencing library plate (2 plates were sequenced per cross, n=96 samples per plate). The black dotted line represents the median read count across all samples (1,802.5), while the red dotted line represents minimal read count threshold for inclusion (251). **b)** Frequencies of the top 2 alleles across all reads in the library. The red line represents the filtering threshold (0.1) under which samples were excluded. **c)** Read count ratio of the second most abundant allele over the first most abundant. The red line represents the threshold (0.4) under which samples were excluded. **d)** Fraction of samples retained after the different filtering steps: minimum read count, top allele frequency and abundance ratio values, and annotation as a pair of LOF mutations based on fitness_IC>100_ scores (see Figure S4).

**Figure S7.**
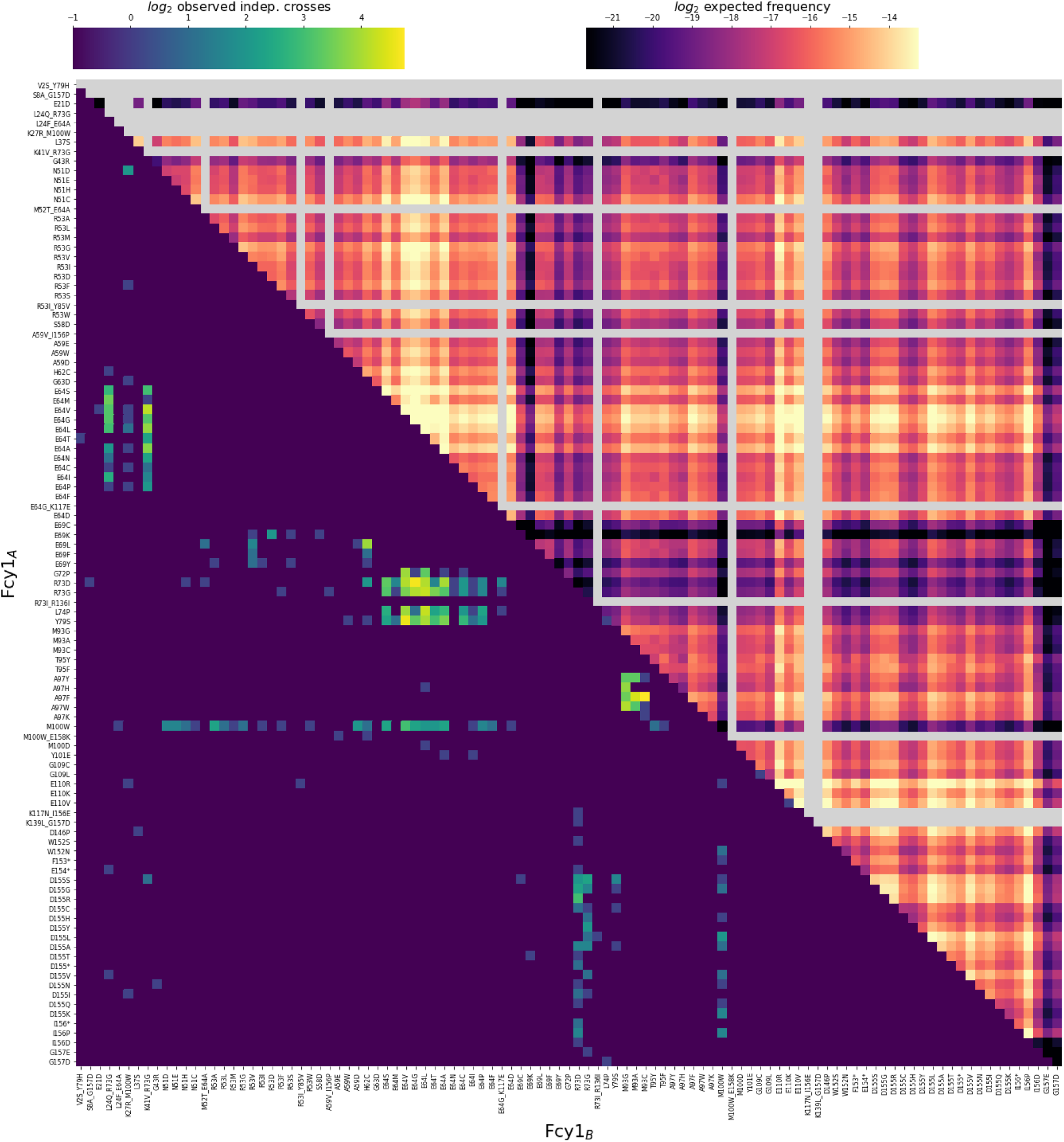
Fcy1 compensating variant pairs identified in the large-scale experiment. The bottom half of the heatmap shows the number of independent crosses observed for each mutant (meaning either the same combination of codons in different mating pools or different combinations of codons within the same mating pools). The top half shows the expected frequency of each mutant pair as expected from the abundance of the two parental alleles in the high-throughput sequencing data.

**Figure S8.**
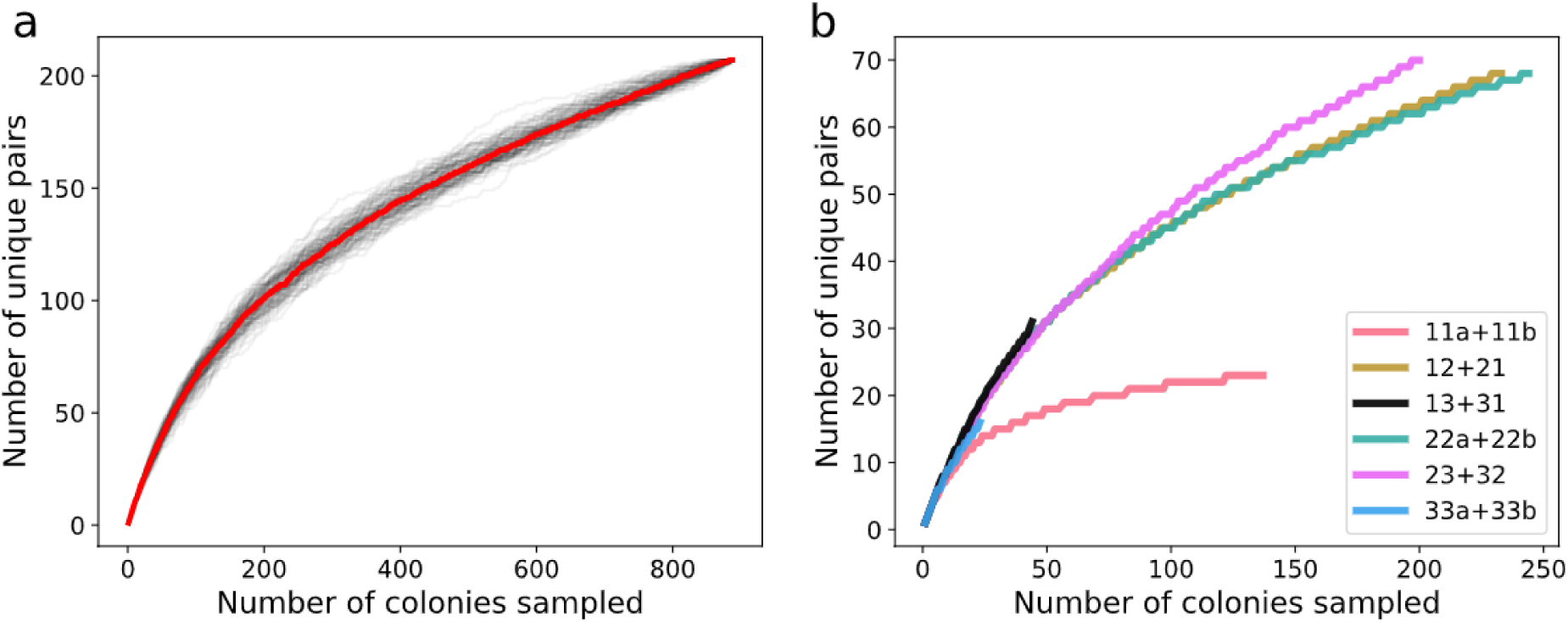
The space of pairs of gene copies showing positive *trans* epistatic is not saturated and varies depending on Fcy1 regions. **a)** Unique gene sequence pairs identified as a function of the number of colonies randomly sampled from our dataset. The red line represents the median for 100 independent sampling iterations, shown as black lines. **b)** Rarefaction curves grouped by cross. Each line is the median of 100 random sampling iterations within both replicate crosses. Fragment 1-Fragment 1 crosses appear saturated in the number of unique pairs observed at a lower level than other variant pool combinations, suggesting less compatible compensating mutations within this region.

**Figure S9:**
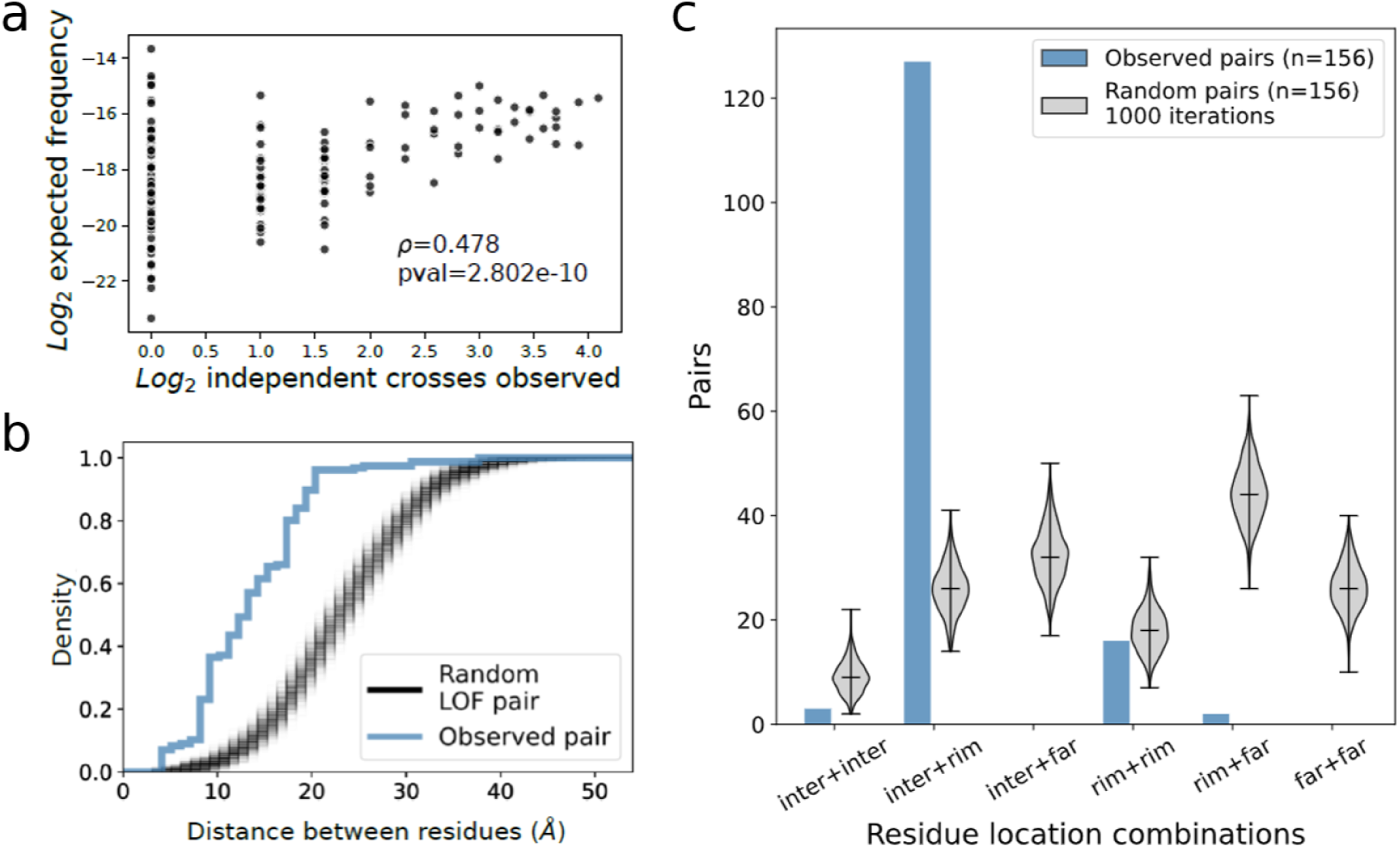
Properties of compensatory variant pairs. **a)** Expected frequency of a mutant pair as a function of the number of independent crossing events sequenced for single mutant pairs (n=156). Spearman’s rank correlation is shown. **b)** Mutant residue to mutant residue distance of screen mutant pairs (n=156 single mutant pairs) compared to random sets of LOF allele pairs (n=156 pairs, 1,000 iterations). Sampling takes into account the distribution of variants across the Fcy1 sequence. **c)** Observed proportion of variant location relative to the interface compared to sets of random LOF allele pairs (inter = at the interface, rim = at interface rim, far = alpha carbon >6 Å from the other subunit). The black bars represent the median and the extremes of the distribution for each combination.

**Figure S10:**
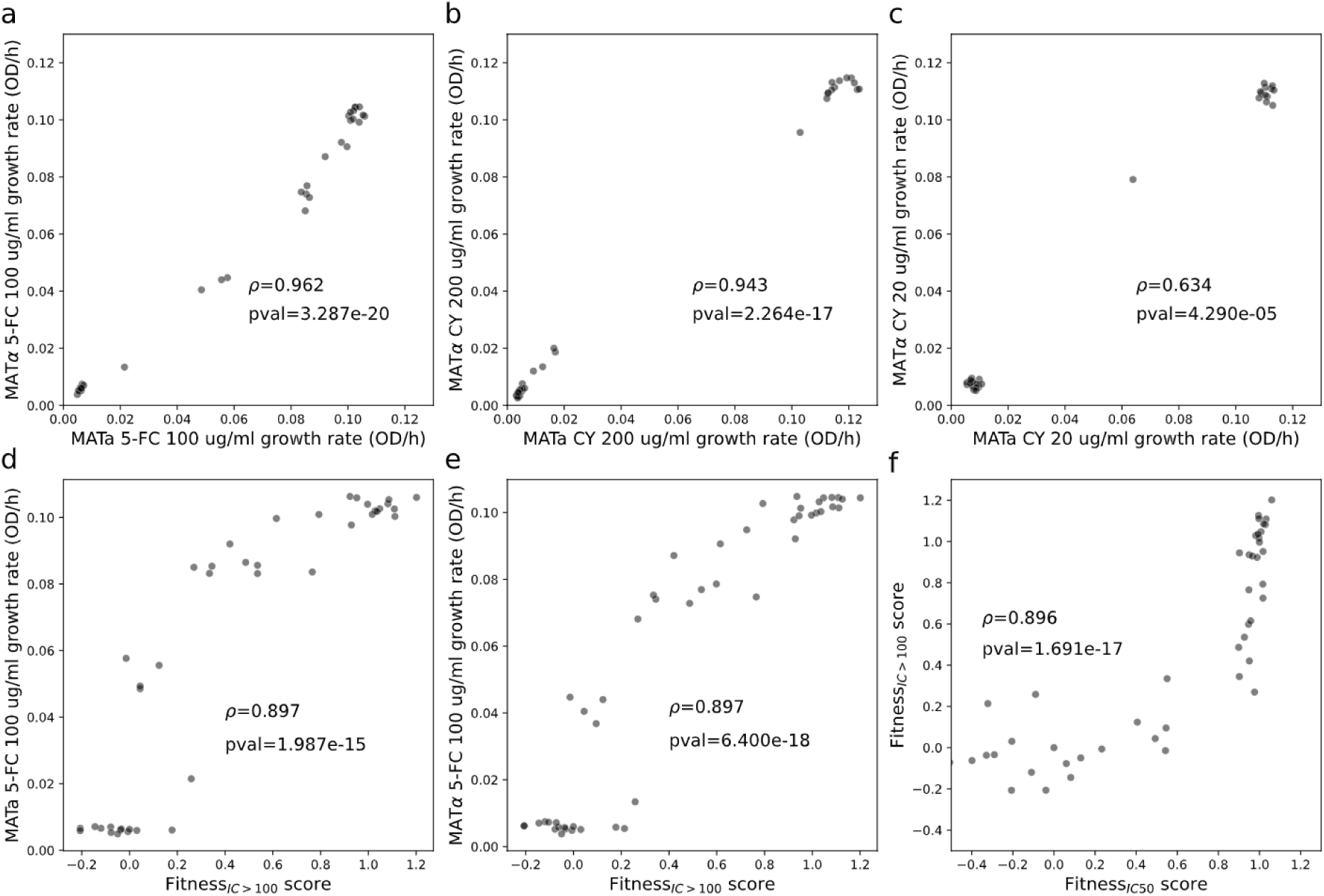
Phenotypes of l FCY1 sequence variants are reproducible across both mating types and are consistent with DMS results. **a)** Growth rate of matched mutants constructed in either MATa or MATα backgrounds in 5-FC_IC>100_ screen conditions (n=35). **b)** and **c)** Growth rate of matched mutants in SC-Ura + 200 μg/mL cytosine (CY) (n=35) and SC-Ura + 20 μg/mL cytosine (n=35). **d)** and **e)** Correlation between MATa (n=42) or MATα (n=48) 100 μg/mL 5-FC growth rate from individual growth experiments and fitness score as assayed in the DMS assay. Wild-type and deletion mutants were assigned fitness scores based on the median score of either synonymous or nonsense mutants in the assay. **f)** Correlation between fitness scores of the DMS assay and fitness scores of (*11*) MATα validation mutants (n=47).

**Figure S11:**
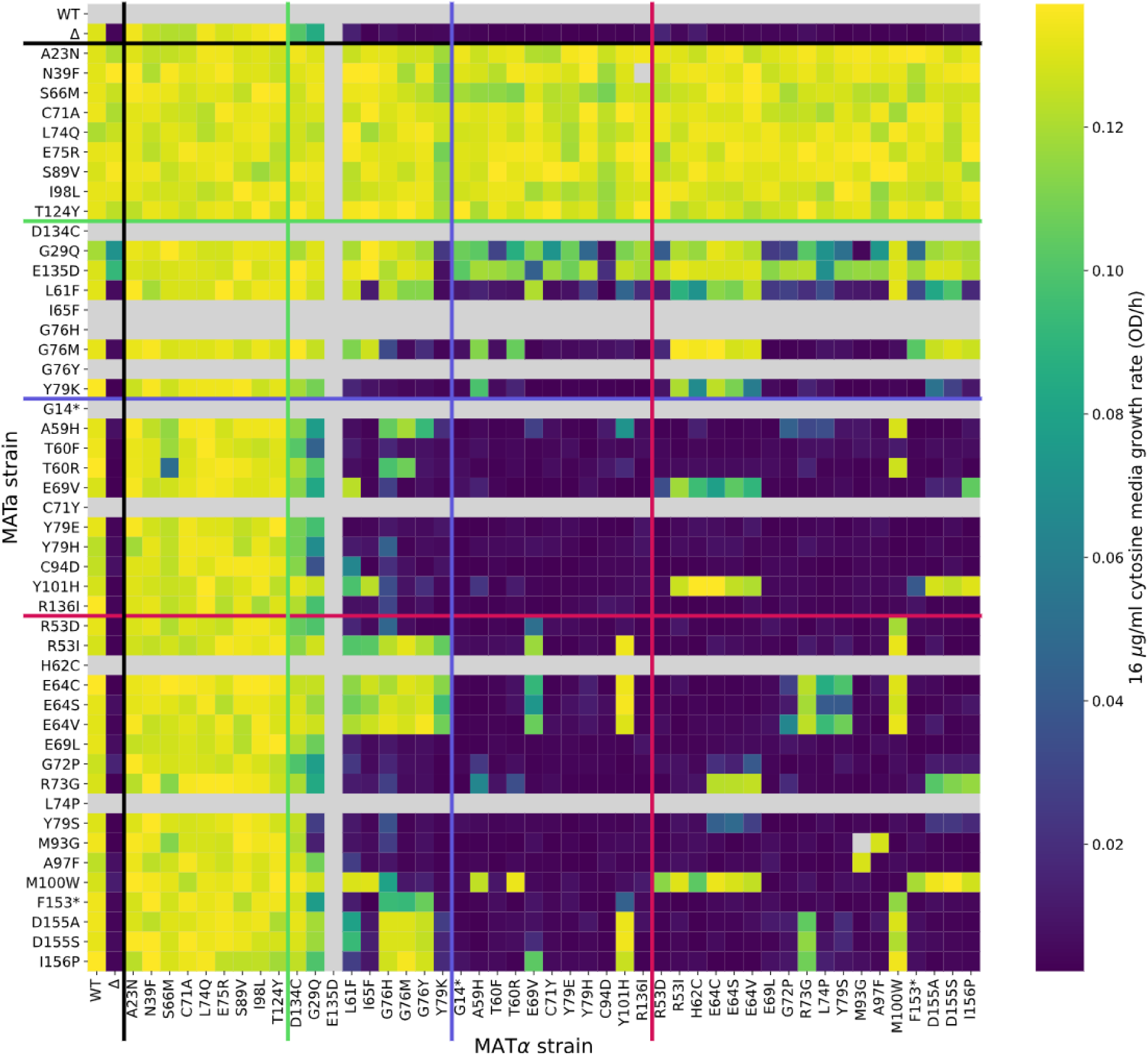
All against all growth assays of FCY1 variant pairs. Coloured lines represent the different groups of mutants that were not observed in the large-scale assay included in the experiment: controls (black), wild-type like (green), mild LOF (blue) and severe LOF (red). Heatmap colour range bounds represent the 2nd and 98th percentiles of the growth rate distribution. The lower right red-bounded square represents the large-scale assay validation subset.

**Figure S12.**
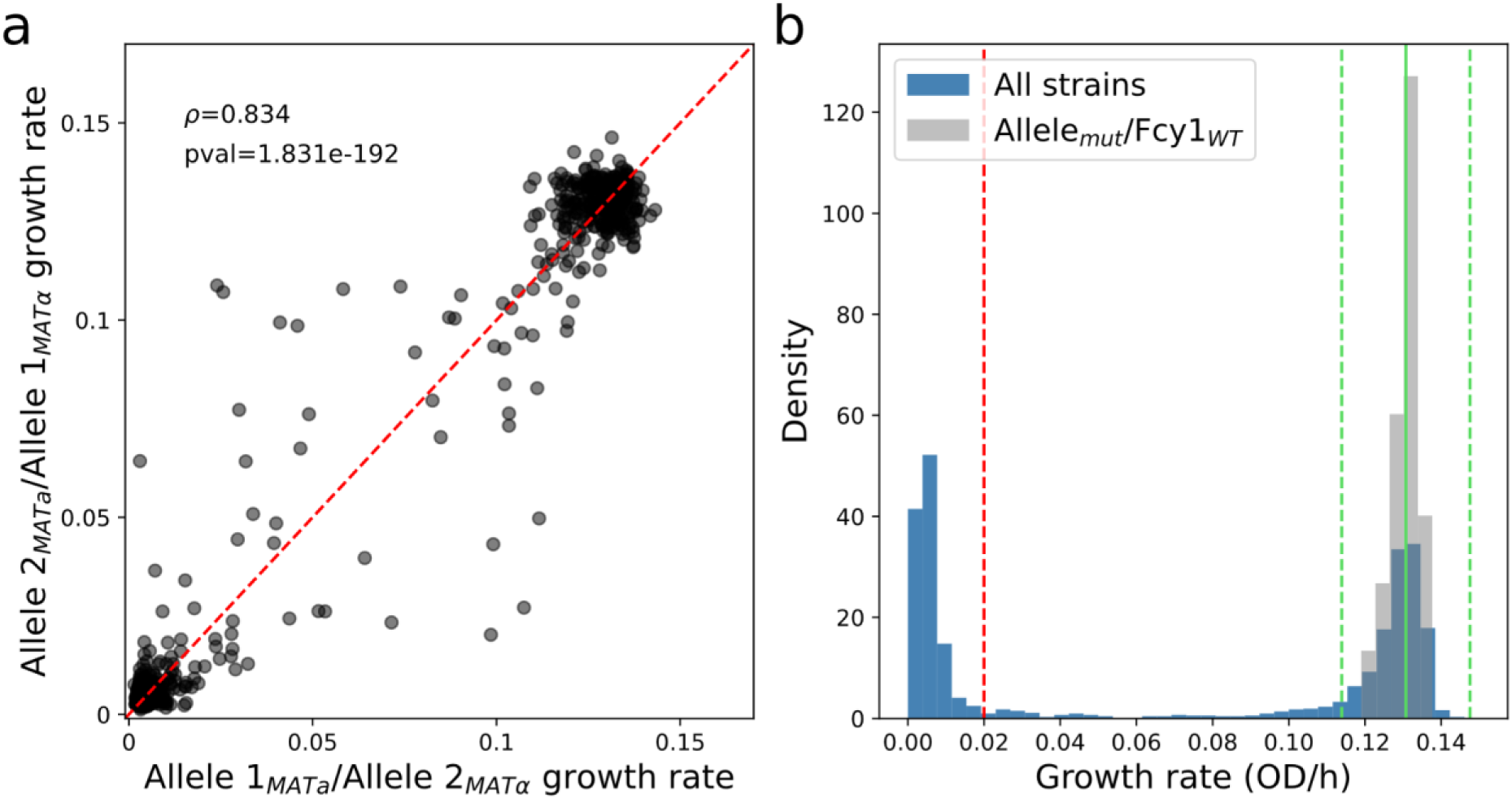
Replicability of validation assay and threshold for compensation event detection. **a)** Correlation between replicate cross maximum growth rate. Spearman’s correlation is shown (n=740). **b)** Measured growth rate distribution in the validation assay. The threshold for growth/no growth classification is shown as a red dotted line. The mean of the Fcy1_mut_/Fcy1_WT_ growth rates is shown as a green line with +/- 4 standard deviations shown as green dotted lines. Pairs of alleles that deviated from their expected fitness by more than 4 standard deviations of were classified as compensating.

**Figure S13.**
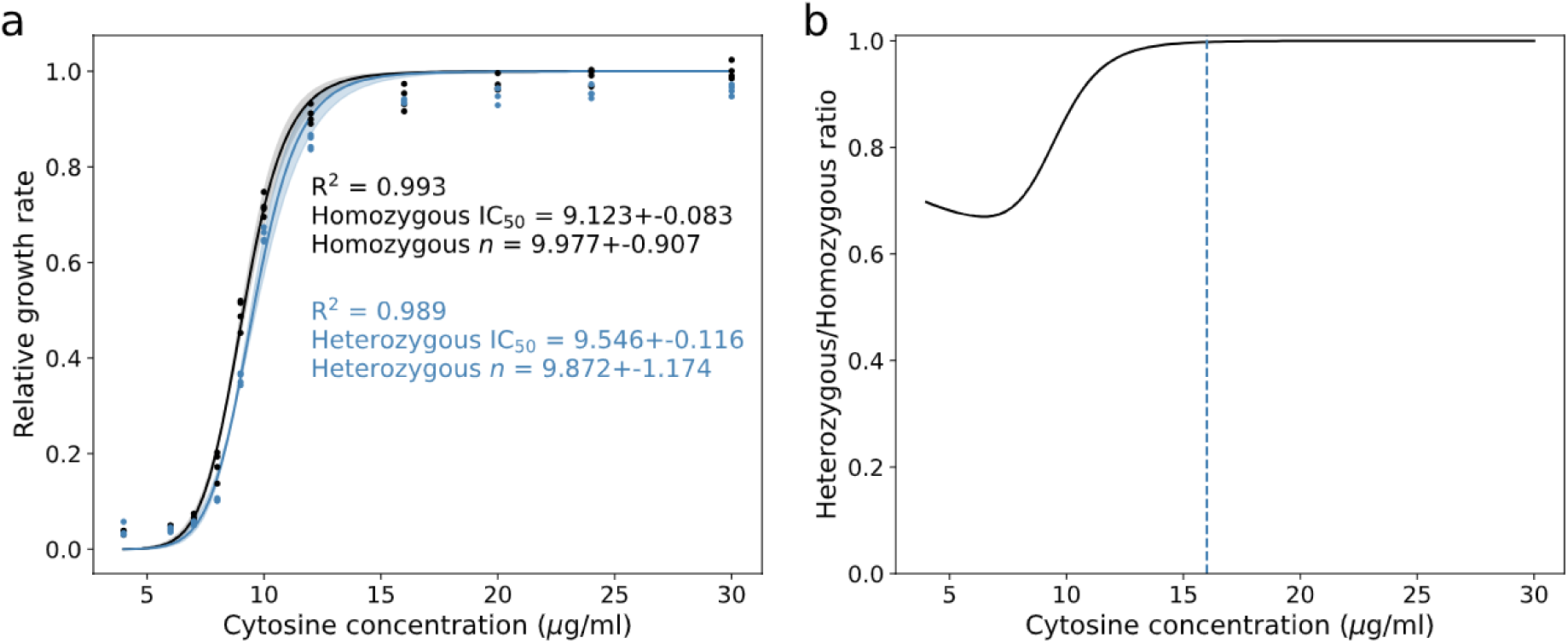
FCY1 is haplosufficient in screen conditions. **a)** Cytosine dose-response curves of homozygous wild-type strains (*FCY1*/*FCY1*) strains and heterozygous deletion strains (two MATa *FCY1/*MATα *fcy1Δ* crosses and two MATa *fcy1Δ//*MATα *FCY1*). Heterozygous deletion strains show a ∼5% difference in the amount of cytosine required to reach 50% growth rate, but no discernible change in sensitivity (*n*). The mean growth rate of homozygous strains at the highest cytosine concentration is used as reference to calculate relative growth rates. **b)** Haploinsufficiency as measured by the ratio between heterozygous and homozygous strains relative growth rates. The dashed line indicates the cytosine concentration used in the large-scale assay.

**Figure S14.**
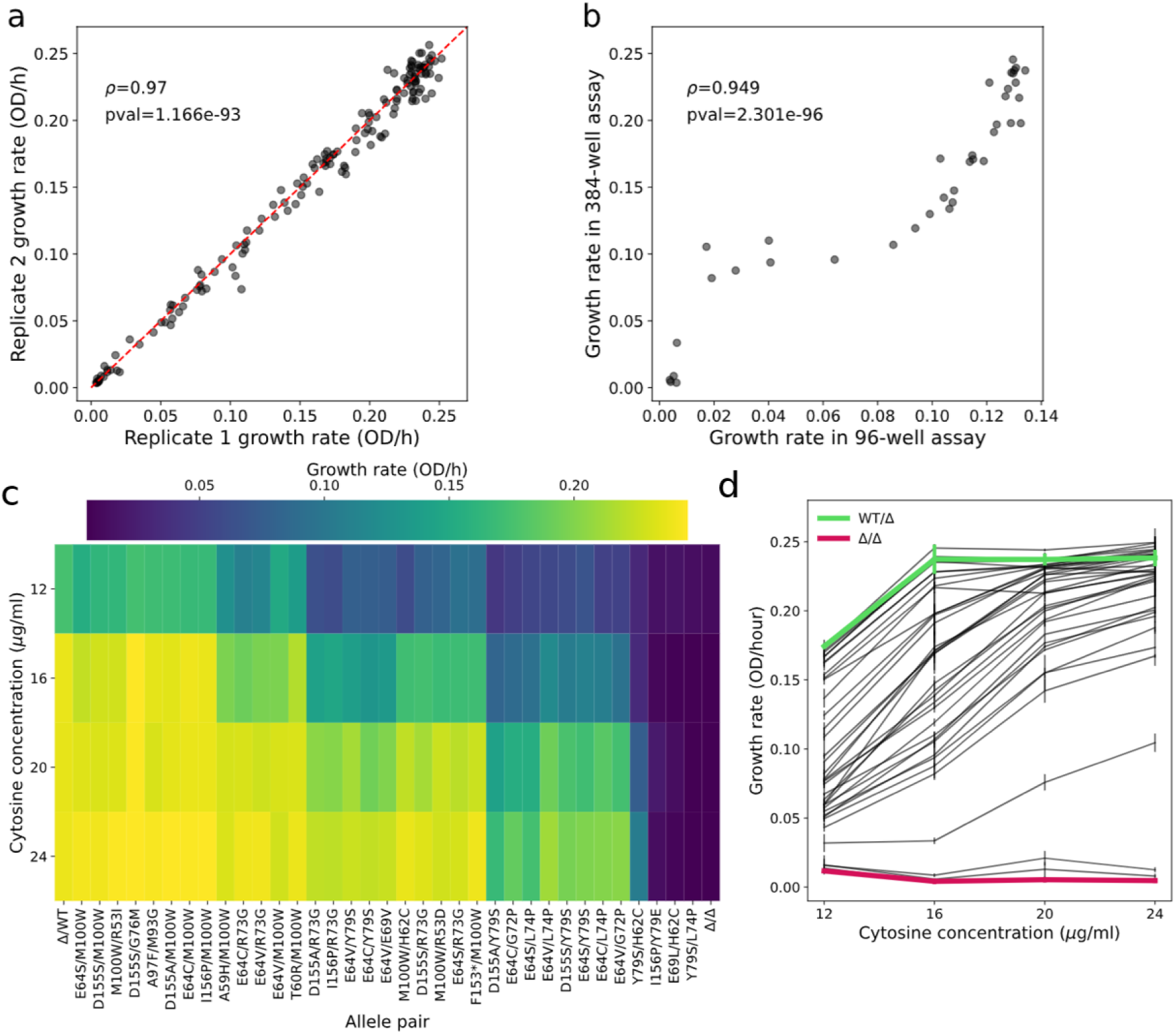
Dose-response curves of compensating mutant pairs. **a)** Correlation between strain by condition replicates in the assay. Spearman’s correlation coefficient is shown (n= 152). **b)** Correlation between strain growth rate in 16 μg/mL cytosine media in the all-by-all validation assay, performed in 96-well plates, and the dose-response assay, performed in a 384-well plate. Spearman’s correlation is shown (n=38). **c)** Growth rate as a function of cytosine concentration for compensating mutant pairs (24 true positives, 3 false positives, and 4 undetected). Heatmap intensity is scaled to the 2nd and 98th percentiles of growth rate values. This assay also shows that one screen hit we failed to validate at 16 μg/ml, H62C/Y79S, can grow at higher cytosine concentrations. Uracil leakage in the large-scale assay might have increased the effective cytosine concentration above 16 μg/ml, explaining this discrepancy (*11*, *57*, *58*). **d)** Dose-response curves of compensating allele pairs compared to the wild-type heterozygous deletion (in green) and the double deletion mutant (in red). Error bars represent one standard deviation around the mean growth rate measurements (n=2).

**Figure S15:**
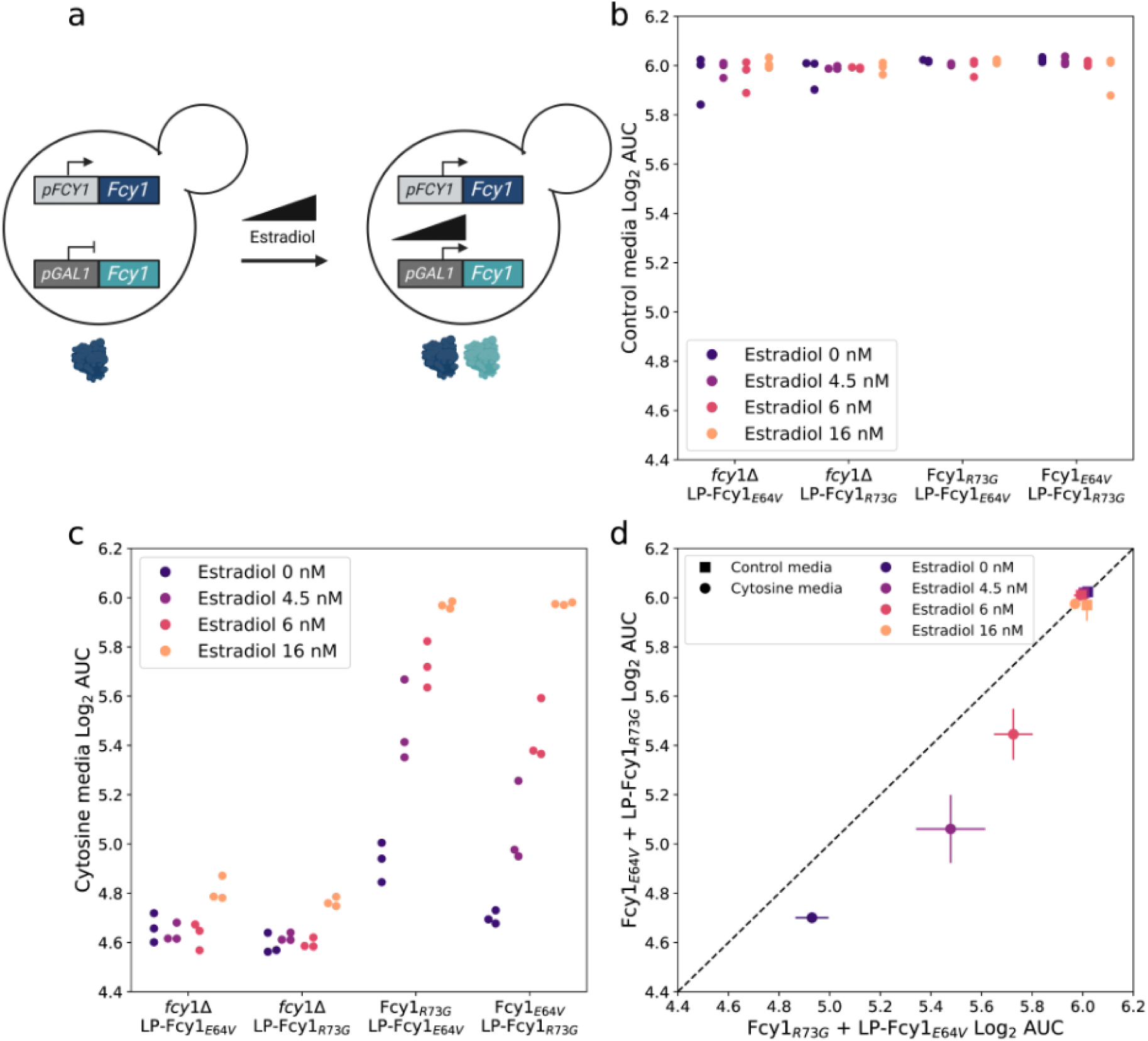
Auxotrophy complementation by Fcy1_E64V_ and Fcy1_R73G_ is expression dependent and does not depend on a diploid background. **a)** Within the same haploid strain, a mutant allele of Fcy1 is integrated at the native locus. Another mutant allele is introduced at genomic Landing Pad, where its expression is under the control of the GAL1 promoter. When estradiol is present in the media, the GEM synthetic transcription factor induces the expression of the LP allele in a dose dependent manner. **b)** Growth of Δ/LP and Fcy1/LP strains SC-leu control media. No difference in fitness is observed between strains. **c)** Growth of Δ/LP and Fcy1/LP strains in modified cytosine media (see methods). **d)** Expression of Fcy1_E64V_ from the inducible Gal1pr locus complements constitutive Fcy1_R73G_ better than the opposite scenario except at maximal induction levels. Panel a was made with Biorender.

**Figure S16.**
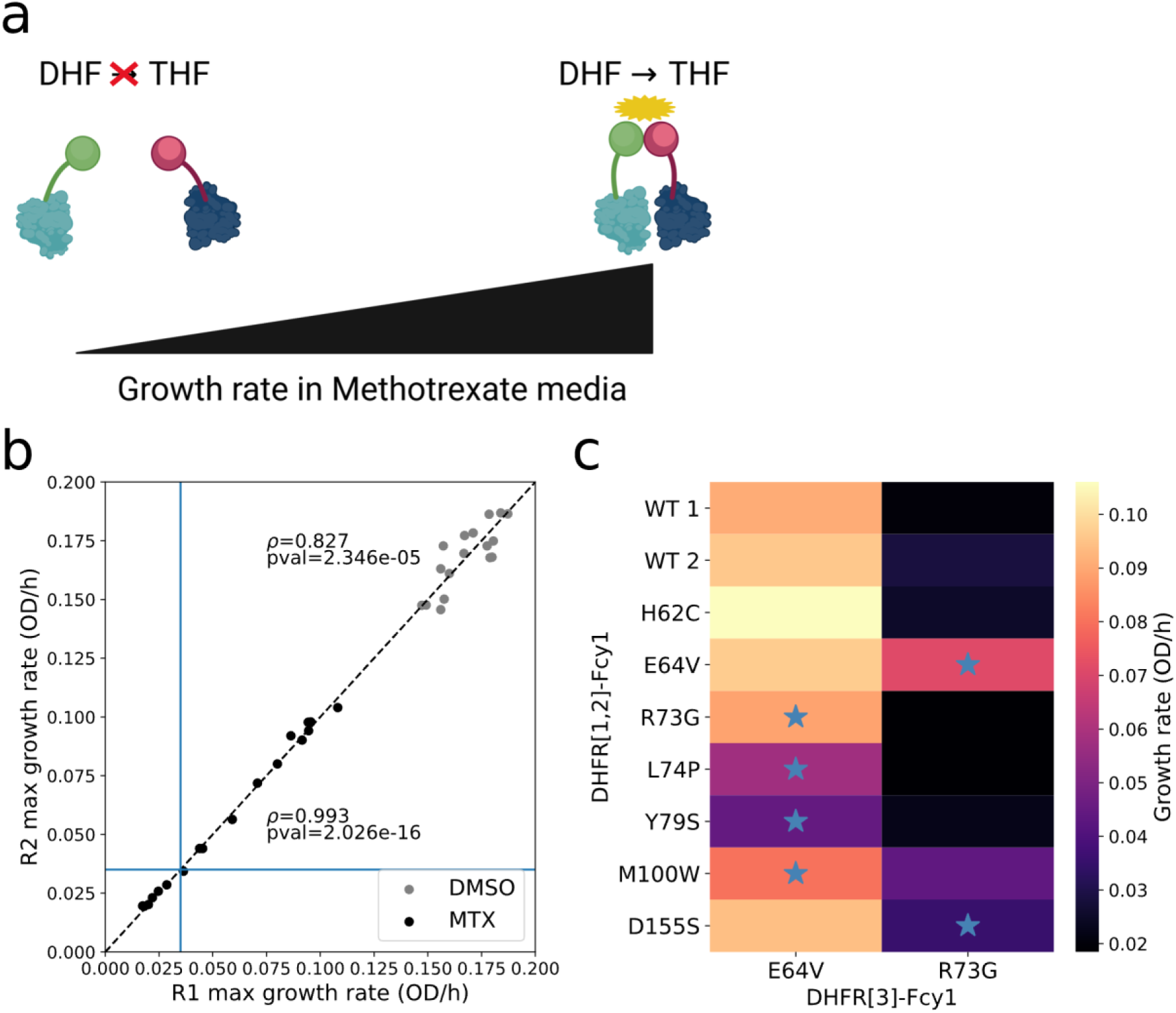
DHFR-PCA detects heteromeric complexes *in vivo*. **a)** Principle of the Dihydrofolate reductase protein fragment assay (DHFR-PCA). The two proteins of interest are tagged with complementary fragments of a methotrexate (MTX) resistant DHFR enzyme. When the two proteins interact together, the fragments assemble into a functional enzyme that allows growth in media where methotrexate inhibits the endogenous DHFR enzyme. Growth is proportional to the amount of interactions between partners, which is a function of the strength of the interaction and the concentration of protein. **b)** Correlation between replicates of each strain in control media (DMSO) or MTX media. Spearman’s correlation coefficient is shown for each condition (n=9). Growth rate in DMSO did not correlate with growth in MTX (ρ=-0.100, pval=0.693). The growth rate cut-off for positive interactions 0.035 is shown as a blue line. **c)** Growth rates of Fcy1 variant pairs in MTX media, which measures protein complex abundance (dependent on both affinity and expression level) *in vivo*. Blue stars denote the six positive *trans* interacting Fcy1 variant pairs covered by this experiment. While the catalytic dead mutant E64V can form complexes with wild-type (WT) Fcy1 and all other variants, the loop mutant R73G appears unable to interact with most variants. This could be explained by its instability, as revealed by other experiments. Panel a was made with Biorender.

**Figure S17.**
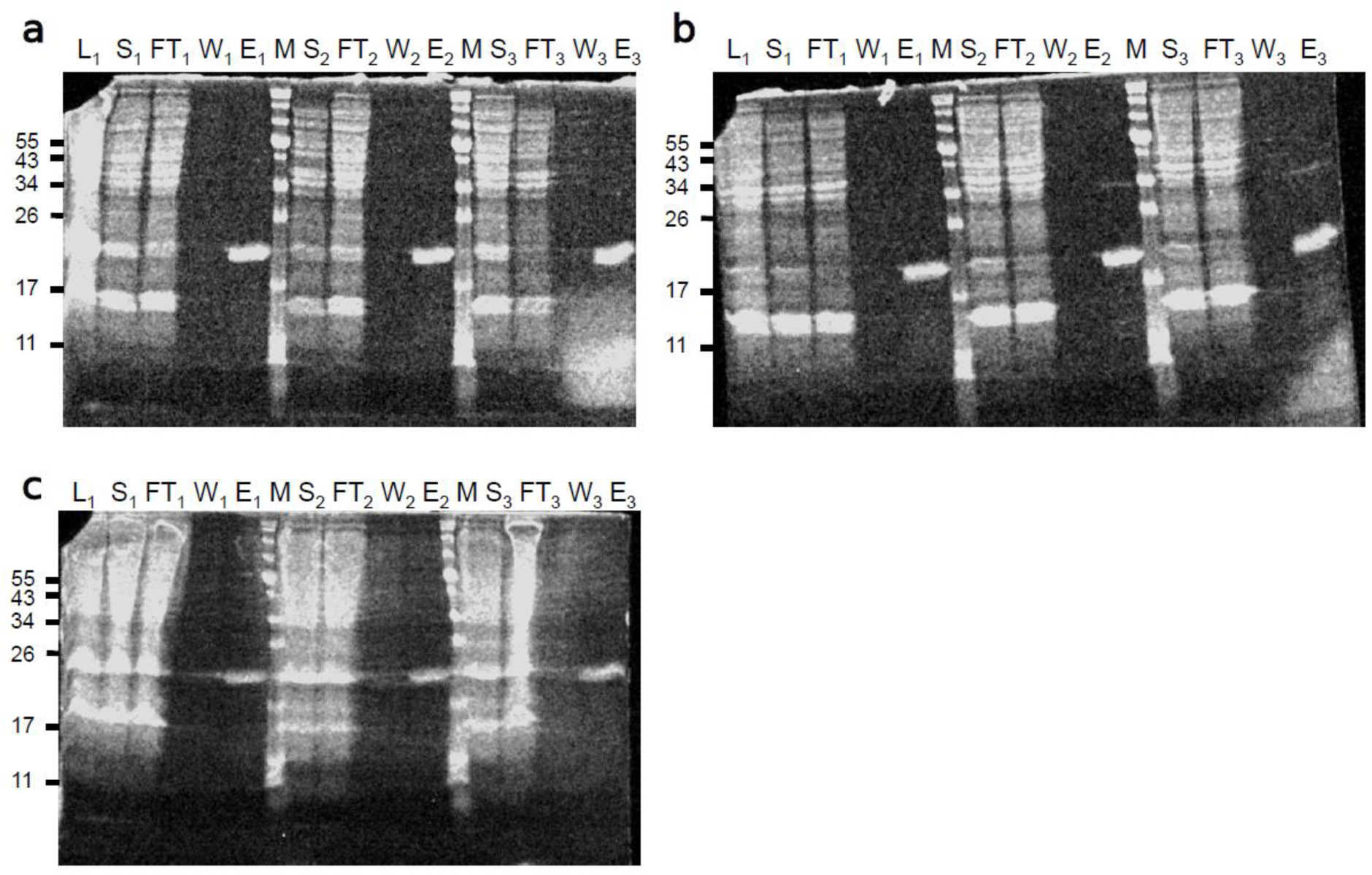
Heterologous expression and purification of Fcy1 and variants yields pure proteins. **a)**, **b)** and **c)** Coomasie stained SDS-PAGE for three affinity purification replicates of Fcy1_WT_, Fcy1_E64V_ and Fcy1_R73G_ using the Dynabeads His-Tag Isolation and Pull-down kit. L: cell lysate, S: soluble fraction, FT: beads flowthrough, W: washes, E: elution, M: NEB Blue Prestained Protein Standard, Broad Range (P7718S). Only one lysate per variant was loaded. For each sample, 25 ul was loaded per well (10 ul sample, 5 ul loading buffer 5X, 2.5 ul DTT 1M, 7.5 ul ddH_2_O) in a 15% acrylamide gel. Gels were run for 55 minutes at 175V and colored 20 minutes in coomassie-R250.

